# A microfluidic rheometer for tumor mechanics and invasion studies

**DOI:** 10.1101/2025.09.29.679368

**Authors:** Young Joon Suh, Manli Liu, Bangguo Zhu, Mrinal Pandey, Brian Cheung, Jaemin Kim, Nikolaos Bouklas, Chris Roh, Jeffrey E Segall, Chung-Yuen Hui, Mingming Wu

**Affiliations:** Department of Biological and Environmental Engineering, Cornell University, Ithaca, NY, USA; Department of Mechanical and Aerospace Engineering, Cornell University, Ithaca, NY, USA; Department of Pathology, Albert Einstein College of Medicine, Bronx, NY, USA

## Abstract

Clinically, the feel, touch, and shape of a solid tumor are important diagnostic methods for determining the malignant state of the disease. However, there are limited tools for quantifying the mechanics and the malignancy of the tumor in a physiologically realistic setting. Here, we developed a microfluidic rheometer – termed the microrheometer – that enables simultaneous measurements of tumor spheroid mechanics and their invasiveness into a 3D extracellular matrix (ECM). The microrheometer consists of a pneumatic pressure control unit for applying controlled static or cyclic compression to tumor spheroids, and a sample chamber for containing spheroid embedded ECM. The innovation here lies in the integration of a polyacrylamide membrane force sensor within the sample chamber, enabling a direct force measurement in a physiologically relevant setting. We found that both tumor stiffness and the viscoelastic properties of the tumor are closely correlated with tumor invasiveness. The microrheometer allowed us to measure tumor mechanics in a short time (less than a minute) and has the potential to be used clinically in the future. We note that the microrheometer here can be easily extended to studies of mechanics of single cell, nucleus, as well as other cell/tissue types.

## Introduction

Mechanical properties of biological tissues play a crucial role in development, wound healing and disease progression ^1^. It is now well accepted that the mechanical properties of extracellular matrices (ECMs) are key determinants of cell growth, differentiation and migration ^2-4^, important for disease diagnosis and prognosis. For example, cancer progression has been linked to increased ECM stiffness, with tumors exhibiting stiffnesses 3-10 times higher than normal tissue ^5-8^. It has been reported that increased tissue stiffness is an important biomarker for breast cancer ^9-12^, liver fibrosis ^13^, pulmonary fibrosis ^14^, sclerosis ^15^, and retinal diseases ^16, 17^. As such, tumor mechanical properties underlie the clinical practice of palpation, where physicians detect hard lumps indicative of pathological changes. However, in early-stage tumors, the stiffness change may be subtle and not reach the 3-10-fold difference, making diagnosing by palpation difficult.

Broadly, tumor mechanics can be investigated under two principal modes of mechanical loading: tension and compression. As tumors grow within confined tissue spaces, compressive stresses accumulate in the tumor core, while tensile stresses accrue at the periphery of the tumor ^18, 19^. Seminal works by the Weaver Lab and others have demonstrated that the tensional state of the ECM promotes tumor malignancy in breast tumor models ^8, 20^. Complementing these findings, work from the Chen Lab has advanced our understanding of cell-generated tensile forces in microtissue physiology ^21^. In contrast, the effects of compressive forces on tumors remain poorly understood mainly due to the lack of appropriate experimental platforms. Current understanding regarding compressive stress has been studied in vitro by growing tumor spheroids within varying concentrations of polymer matrices such as agarose ^22-24^. These studies reveal the importance of compressive stress in tumor progression. For recapitulating a physiologically realistic tumor microenvironment, where compressive stress is anisotropic and dynamic, it is important to develop tools that can apply controlled uniaxial and dynamic compressions to microtissues in a 3D environment.

The main technique for the mechanical characterization of soft materials is the commercially available dynamic rheometer. Typically, a tissue of certain geometry (e.g. a cylinder) is mounted between two parallel plates, and a sinusoidal torsional shear or compression force is applied to the tissue, and the stress-strain curve is used to infer the material properties. This measurement provides precise viscoelastic properties of soft materials and is straightforward to use ^25^. It has provided much of what we know about the mechanical properties of biomaterials today ^8, 25^. However, its application to living tissues is limited due to its incompatibility with tissue culture conditions and real-time microscopic imaging. Atomic Force Microscopy (AFM), a widely used technique for measuring cell and tissue mechanics, enables simultaneous mechanical characterization and imaging of single-cell dynamics ^26-28^. It provides valuable insight into localized tissue mechanics with high spatial resolution ^8^. However, a major limitation of AFM is its low throughput, primarily due to the spatial heterogeneity of cells and tissues, which require point-by-point measurements across the sample. Other microrheology techniques, including optical tweezers ^29, 30^ and magnetic bead-based methods ^31-35^, allow for precise force application in a targeted position, but they often suffer from limitations in force magnitude or the depth of field for tissue samples. Micropipette aspiration, which measures tissue deformation under suction pressure, is relatively straightforward to use for single-cell mechanical measurements, but it lacks compatibility with high-throughput or integrated live-cell imaging systems ^36, 37^.

Microfluidic technologies have emerged as effective platforms for probing single-cell/tissue mechanics while maintaining compatibility with live microscopic imaging. Microfluidic constriction devices have been used to derive viscoelastic properties of cells, including elastic modulus and fluidity, from dynamic shape changes during transit through the constriction passages ^38-41^. Similarly, flow-based microfluidic devices have been developed to indirectly measure the tissue mechanics by analysing deformation under controlled shear flow conditions ^42-44^. A major advantage of these systems is their high-throughput capability, enabling rapid mechanical assessment across large populations of cells. In contrast to these flow-based systems, where the cells were suspended in fluid, microfluidic compression platforms have been developed to apply direct compression to single cells ^45-57^ and cell aggregates ^58^. However, most of these platforms lack integrated force sensing, making them unsuitable for direct modulus measurement. While these platforms have greatly advanced the study of single-cell or tissue mechanics, they are typically limited to 2D or suspended cell environments and do not replicate physiological conditions such as a 3D ECM.

In this manuscript, we present a microfluidic rheometer, microrheometer, that introduced a novel in situ force sensor enabling a simultaneous investigation of tumor tissue mechanics and single-cell dynamics in a physiologically realistic 3D setting.

## Experimental methods

### Cells, spheroids, and 3D culture

#### Cells

Two breast tumor cell lines with known malignant levels and surface adhesion molecules (E-cadherin and integrin) were chosen for this work ^59^. MDA-MB-231 is a metastatic triple-negative breast tumor cell line. It was obtained from ATCC (Cat. #: HTB-26, ATCC, Manassas, VA) and cultured in high glucose Dulbecco’s modified eagle medium (DMEM) (Cat. #: 11965092, Gibco, Life Technologies Corporation, Grand Island, NY) supplemented with 10% fetal bovine serum (Cat. #: S11150, Atlanta Biologicals Lawrenceville, GA), 100 units/mL penicillin, and 100 μg/mL streptomycin (Cat. #: 15140122, Gibco). MCF-10A, a non-tumorigenic breast epithelial cell line serving as a control, was obtained from ATCC and cultured in DMEM/F-12 media (Cat. #: 11320033, Gibco) supplemented with 5% horse serum (Cat. #: S12150, Gibco), 20 ng/mL hEGF (Cat #: PHG0311, Gibco), 0.5 µg/mL hydrocortisone (Cat. #: H0888-1G, Sigma-Aldrich), 100 ng/mL cholera toxin (Cat #: C-8052, Sigma), 10 μg/mL insulin (Cat. #: I-1882, Sigma), 100 units/ mL penicillin, and 100 μg/mL streptomycin (Cat. #: 15140122, Invitrogen). All cells were cultured in T75 flasks (Cat. #: 10062-860, Corning, Lowell, MA, USA), which were placed in an incubator set to 5% CO_2_, 37°C, and 100% humidity. Cells were passaged every 3–4 days and harvested for experiments when the cell culture reached 70–90% confluency. MDA-MB-231 and MCF-10A cells with 20 or fewer passages after acquisition from ATCC were used in all experiments.

#### Spheroids

Uniformly sized tumor spheroids were formed using a specially designed microwell array (Fig. S1) as described in ref. ^60, 61^. Briefly, an array of 36 x 36 microwells was first patterned on a 1 mm-thick agarose gel membrane using a soft lithography method. Each microwell is cylindrical in shape with a diameter of 200 µm and a depth of 250 µm. The agarose gel surface provides low adhesion surfaces to the cells, making it easier for the cells to cluster together and form spheroids. One microwell array was then placed in each well of a 12-well plate (Cat. #: 07-200-82, Corning, Lowell, MA, USA). Within each well, we placed 2 million MDA-MB-231 cells or 3 million MCF-10A cells suspended in 3 mL of the same media as the one we used for MCF-10A cell culture. The 12-well plate was then kept in an incubator (Forma, Thermo Scientific, Asheville, NC, USA) at 37 °C, 5% carbon dioxide, and 100% humidity for 7 days before harvesting. On days 3 and 6, the medium was changed to fresh medium. The average diameter of the spheroids was about 130 *μ*m at the time of collection. For each experiment, the spheroids were collected from one microwell array, and a Falcon^®^ Cell Strainer (Cat. #: 10054-458, Corning, Lowell, MA, USA) with 100 μm pores was used to collect the spheroids. We note that MCF-10A cells form spheroids primarily through direct cell-cell adhesion mediated by E-cadherin, whereas MDA-MB-231 cells form spheroids indirectly via cell-ECM adhesion through integrins ^62^. For this reason, rich media (DMEM/F12) and long culture time (7 days) are important for the formation of MDA-MB-231 spheroids.

#### 3D spheroid culture

Tumor spheroids were embedded within 3.5 mg/mL type I collagen. To make 3D spheroid culture, first, a 68.16 μL collagen stock (10.27 mg/mL, Cat. #: 354249, Corning, Lowell, MA, USA) was titrated with 1.5 μL 1 N NaOH. Second, 20 μL 10X M199 (Cat. #: M0650-100ML, Sigma) was added to approximately yield a final pH of 7.4. Third, 110.34 μL of spheroids suspended in cell media was added to reach a final volume of 200 μL and a final collagen and spheroid concentration of 3.5 mg/mL and 2860 spheroids/mL, respectively.

### Fabrication of the microrheometer

The microrheometer consists of three layers, all of which are made of PDMS and casted from Silicon master molds. The silicon masters are fabricated using photolithography techniques at the Cornell Nanoscale Science and Technology Facility.

#### Silicon Master Fabrication

The silicon master for the sample chamber layer (L1) is a 2 x 6 array of cylindrical chambers with a rectangular wall at the periphery (Fig. S2A). To ensure the uniform thickness across all the sample chamber layer (L1), SUEX film (Cat. #: SUEX K500 and K200, DJ Microlaminates, Sudbury, MA) of various thickness (e.g. 500 µm and 200 µm) were used. To fabricate the master for L1, first, 100 mm single side polished (P or N type) silicon wafers were cleaned with the piranha solution and dehydrated in a 90^°^C oven overnight. The wafer was then treated with a high-power oxygen plasma (100 W) for 2 minutes in a Glen 1000 plasma cleaner to enhance the bonding strength to the SUEX film. Second, a 500 µm thick SUEX film was laminated on the silicon wafer at 75^°^C at a speed of 1 ft/min using a laminator (Cat #: SKY-335R6, Seoul, Korea). For the L1 with 700 µm thickness, an additional 200 µm thick SUEX film was laminated on top of the 500 µm SUEX film. Third, the SUEX film was exposed to UV light (365 nm) through a pre-patterned mask using an ABM contact aligner, with exposure doses of 2500 mJ/cm^2^ for the 500 µm film and 4500 mJ/cm^2^ for the 700 µm film. The SUEX film was then post-exposure baked at 65^°^C for 15 minutes and then developed in the EBR-10A (PGMEA) developer for 1 hour with no agitation. Finally, the wafer was coated with 1H,1H,2H,2H-Perfluorooctyltriethoxysilane (FOTS) using the MVD100 to facilitate PDMS detachment in later steps.

The silicon master for the deformable piston membrane layer (L2) is a 2 x 6 array of 300 µm cylindrical wells (Fig. S2B). To fabricate the silicon master for L2, 300 µm cylindrical wells were etched into the silicon wafer. Briefly, 4.5 µm thick SPR-220-4.5 photoresist was spun on a clean and dehydrated 100 mm Silicon wafer. The photoresist was then baked at 115^°^C for 2 minutes on a proximity hot plate. Then, it was exposed under a piston-patterned mask with exposure dose of 120 mJ/cm^2^ using an ABM contact aligner. After allowing the post-exposure reaction to take place at room temperature for 30 minutes, it was then developed in the AZ 726 MIF developer for 120 seconds. Then, a mild descum process was completed using an Oxford 81 etcher for 90 seconds. Finally, the Silicon wafer was loaded on the Unaxis 770 Deep Si Etcher, and a total of 567 (200 + 200 + 167) loops of Bosch process was performed to etch 300 µm into the Silicon wafer. To remove the remaining photoresist, the wafer was exposed to a strong plasma in a EcoClean Asher. The wafer was then coated with FOTS using the MVD100. The depth of the piston wells was carefully measured using the P-7 Profilometer throughout the process to confirm the depth.

The silicon master for the pressure control chamber layer (L3) is two parallel channels with three cylindrical chambers with a diameter of 3 mm with an inlet (Fig. S2C). To fabricate the silicon master for L3, SU-8 Epoxy photoresist was used. After cleaning and dehydrating a 100 mm Silicon wafer, SU-8 100 photoresist was spun at 1300 rpm for 30 seconds with a ramp rate of 300 rpm/s. The wafer was soft baked at 65^°^C for 30 minutes and then at 95^°^C for 24 hours. The wafer was then cooled down to room temperature at a rate of 1^°^C/min. The SU-8 photoresist was then exposed under a patterned mask at a dose of 650 mJ/cm^2^ using an ABM contact aligner. Then, the wafer was post-exposure baked at 65^°^C for 5 minutes and at 95^°^C for 30 minutes. The wafer was then cooled down to room temperature at a rate of 1^°^C/min. The SU-8 photoresist was then developed in the SU-8 developer solution with weak agitation for 1 hour. Finally, the wafer was then coated with FOTS via molecular vapor deposition in the MVD100.

#### Soft lithography

All three layers of the microrheometer were fabricated using soft lithography techniques using the silicon masters The sample chamber layer L1 is made via a PDMS double casting method. To make the first PDMS mold, 25 g of 10:1 polydimethylsiloxane (PDMS) mixture was prepared, degassed, and poured into a 5.3 mm thick polycarbonate frame placed around the feature on the silicon wafer. A sheet of laser printer transparency film was placed on top to create a smooth surface, followed by a glass panel with a weight to ensure even thickness. The PDMS was then cured at 60^°^C overnight. The cured PDMS was cut, removed from the silicon master, plasma cleaned for 1 minute, and treated with a homemade FOTS setup (Fig. S3) overnight. This PDMS mold is now ready for the second casting. To cast the sample chamber layer L1 from the PDMS mold, another 15 g of 10:1 PDMS was prepared and degassed. A clean 25 mm x 75 mm x 1 mm glass slide was taped on all four sides onto the bottom of a petri dish, and PDMS was poured on top of the glass slide and the feature side of the PDMS mold. After removing all bubbles in a vacuum chamber, the PDMS mold was flipped onto the slide, secured with tape, weighted down flush such that the height of the device is defined by the PDMS mold. The assembly was cured overnight at 60^°^C and then carefully peeled away, leaving the L1 securely attached to the glass slide.

To fabricate the piston membrane layer L2, 10 g of 10:1 PDMS was prepared and degassed .A 340 µm thick rectangular frame was placed around the feature on the Silicon wafer, and PDMS was poured into the frame. A sheet of transparency film was carefully placed on top without introducing bubbles, followed by a glass slide and a weight. The PDMS was cured at 60^°^C overnight. After curing, the frame and excess PDMS were carefully removed, leaving only the center PDMS piece on the wafer ready to be bonded with the L3 later. Later, the height of the piston was confirmed to be 300 µm with a laser scanning profilometer (Fig. S4)

To fabricate the pressure chamber layer L3, 25 g of 10:1 PDMS was prepared and degassed A 5.3 mm thick rectangular frame was positioned on the Silicon wafer, and PDMS was poured into it. A sheet of transparency film was placed on top without introducing bubbles, followed by a glass panel and a weight. The PDMS was cured at 60^°^C overnight. After curing, the PDMS was cut and removed from the wafer, and a 1.5 mm hole was punched at the inlet using a biopsy punch for pressure delivery.

#### Fabrication of an In Situ Force Sensor - Polyacrylamide (PAA) gel membrane formation

A 200 µm thick PAA gel was polymerized at the bottom of each of the 12 sample chambers to serve as a force sensor. To prepare a polyacrylamide gel with a final concentration of 4% Acrylamide and 0.1% Bis, a precursor solution was first prepared from 20 µL of 40% Acrylamide (Cat. #: 161-0140, Bio-Rad, Hercules, California), 10 µL of 2% Bis (Cat. #: 161-0142, Bio-Rad, Hercules, California) solutions, and 150 µL of sterile dH_2_O in an Eppendorf tube and mixed well. The precursor solution was then degassed in a vacuum chamber for 15 minutes. Then, 20 µL of 2.5% w/v LAP (Lithium phenyl-2,4,6-trimethylbenzoylphosphinate) (Cat. #: 900889-1G, Sigma) solution was added to the precursor solution to reach a final volume of 200 µL. Then, nitrogen gas was gently injected into the Eppendorf tube before it was placed in a homemade nitrogen glove box. The glove box was then purged with nitrogen gas until the oxygen concentration reached 0.1% to ensure proper polymerization, since oxygen inhibits the free radical cross-linking of acrylamide monomers. Inside the nitrogen glove box, 5 µL of PAA gel solution was loaded into each sample chamber. To precisely control gel height, a cylindrical microfabricated button with a height of 500 µm and radius of 1500 µm was lowered on to the L1 layer. The PAA gel was then polymerized under 365 nm UV light for 15 minutes. Due to the thin geometry and small volume, the gel was highly susceptible to oxygen diffusion, making strict oxygen control critical for successful polymerization. After polymerization, 2 µL of 0.002% solids 1 µm carboxylate-modified green fluorescent beads (F8823, Thermo Scientific, Asheville, NC, USA) were added to each well and incubated in a 100% humidity chamber for 30 minutes to allow the beads to settle onto the gel surface. The gel was then gently air-dried to promote bead adhesion to the surface of the PAA gel. The device was then immersed in PBS overnight to fully rehydrate the gel. The PAA gel was washed three times with PBS before experiment.

### Microrheometer assembly and operation

#### Pressure control unit assembly

The pressure control unit is made by bonding L2 (deformable PDMS membrane) and L3 (pressure control chamber). Both L2 (attached to the wafer) and L3 PDMS were plasma cleaned for 1 minute in a plasma cleaner (PDC-001-HP, Harrick Plasma, Ithaca, NY, USA). The features were then carefully aligned under a microscope and bonded. The bonded layers were placed in a 90^°^C oven for 15 minutes to strengthen the bond, then cooled slowly to room temperature. The bonded PDMS piece was carefully detached from the wafer and cut to match the size of L2. Lastly, 4 access ports (pink in Fig. 1A) were made using 2 mm biopsy punches to allow media replenishment and gas exchange from outside to the cell chamber.

**Fig. 1.**
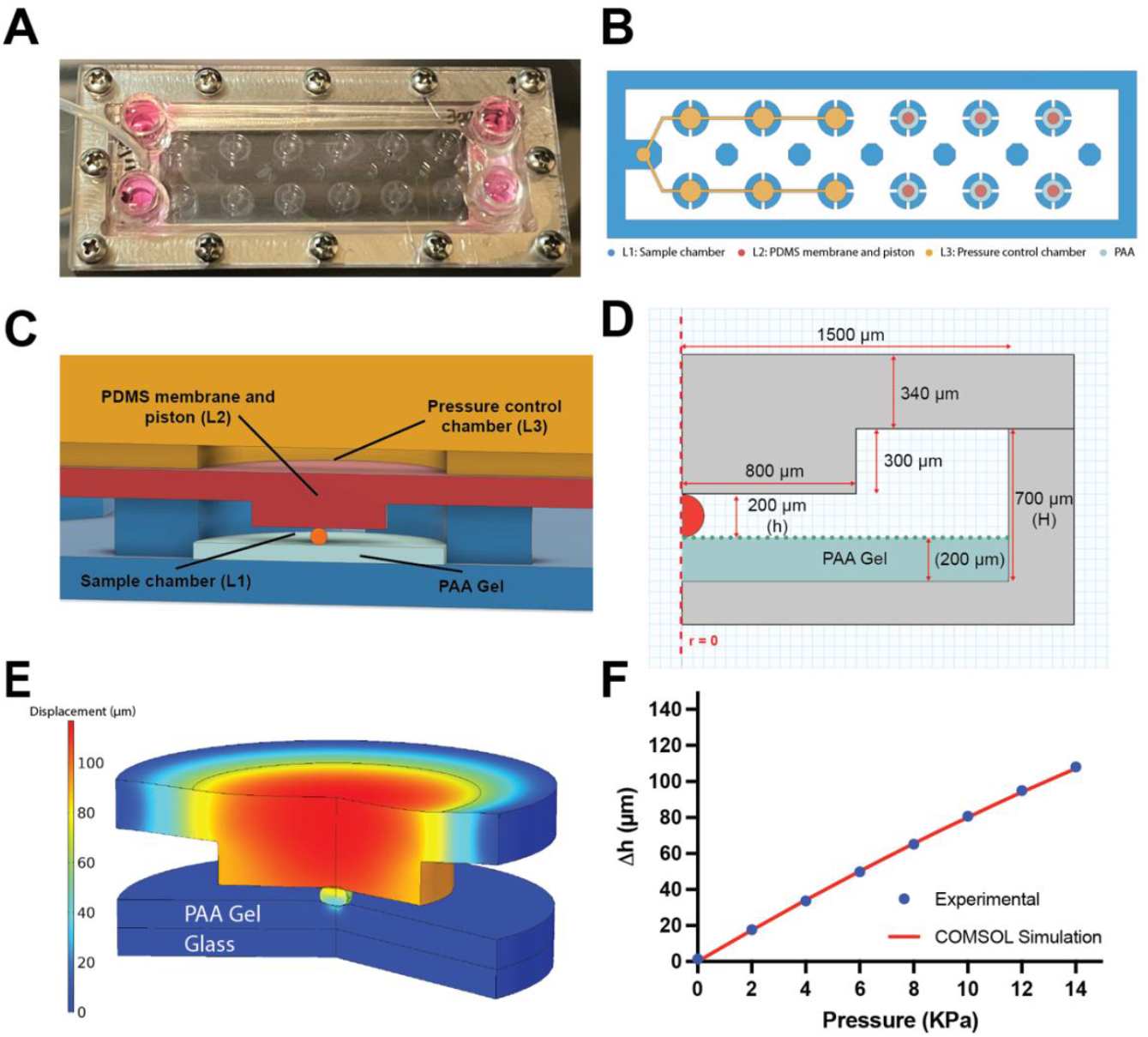
Microrheometer design and calibration. A. A photo of the assembled device. The PDMS device is sandwiched between a stainless-steel frame and a Plexiglas manifold. B. Schematics of the microrheometer. The microrheometer consists of 3 layers: L1 Sample chamber layer (blue), L2 Deformable PDMS membrane and piston layer (red), L3 Pressure control chamber layer (yellow). C. Cross-sectional view of one compression chamber. The pressure in the pressure chamber deflects the membrane such that the piston compresses the tumor spheroid in the sample chamber. D. The dimensions of an axisymmetric compression unit. The dashed line represents the central axis of the compression unit. A critical dimension is h, the distance from the bottom of the piston to the surface of the PAA gel. The sample chamber height is H with a radius of 1500 µm. The thickness of the PDMS deformable membrane is 340 µm, and the height of the piston is 300 µm with a radius of 800 µm. The PAA gel thickness is 200 µm. h and H can be varied depending on the experiments to account for different sample sizes. E. Vertical displacement of the deformable membrane and the piston, spheroid, and PAA gel using COMSOL with pressure of 14 kPa in the pressure control chamber and the dimensions shown in D. F. Calibration curve of the piston displacement (Δh) vs pressure applied in the pressure control chamber. Dots are experimental results, and solid line is the result of COMSOL calculation. Note: Error bars represent standard error of mean (SEM) from 8 experimental values but are too small to be visible due to minimal variation.

#### Microrheometer assembly

Prior to experiment, the pressure control chambers were filled with water to be used for fluid-based pressure control. The surface of the sample loading chamber was treated for better adhesion with the PAA gel or the type 1 collagen gel. For the pressure control unit preparation, the pressure control unit was plasma cleaned for 1 minute and submerged under water overnight to fill the pressure control chamber with water. Then, a tubing (OD and ID of 1/16” and 1/32”, respectively) pre-filled with water from the pressure controller (Elveflow OB1 MK4, Paris, France) was connected to the 1.5 mm inlet on the device. To prepare the sample loading chamber layer L1, each well in the L1 was selectively plasma cleaned for 15 seconds with a mask, treated with 1% Polyethylenimine (PEI) for 10 minutes, washed with sterile deionized water once, and treated with 0.5% glutaraldehyde for 30 minutes. After three washes with sterile deionized water, the L1 was ready for use in the experiment.

The samples were loaded differently depending on the experiment. For tumor mechanics measurement, the tumor spheroids were directly loaded into the sample loading chamber without collagen. For the tumor invasion studies, the spheroid-embedded collagen mixture was loaded into the well. More specifically, 5 µL of spheroid/collagen mixture was loaded into each well of the sample chamber in the L1. The pressure control unit was then aligned under a microscope and lowered onto the L1. The three layers were then sandwiched between a metal frame and a Plexiglas window, secured with screws to ensure uniform pressure around the entire assembly as shown in Fig. 1A. If the sample included collagen, the device was placed in a 37^°^C and 5% CO_2_ incubator for 45 minutes for collagen polymerization. After the polymerization, the device was carefully filled with cell media through the four access ports. Lastly, the device was connected to the Elveflow pressure controller and placed on a temperature, humidity, and CO_2_ controlled microscope stage for imaging.

#### Microrheometer operation

The pressure in the pressure control unit was regulated using a commercially available pressure controller (OB1 MK4, Elveflow Inc., Paris, France), operated via Elvesys software. The software allows users to generate pressure waveforms with customizable profiles. The piston’s stability and responsiveness under pressure inputs were validated (SMovie 1-3). An external pressure source (Model 3 Compressor, Jun-Air, MI, USA) supplied pressure to the OB1 MK4 controller.

### Imaging and data analysis

#### Imaging

All images were taken with a 20x magnification objective lens (NA = 0.25, Olympus America, Center Valley, PA, USA) on an inverted epi-fluorescent microscope (IX 81, Olympus America, Center Valley, PA, USA) and a CCD camera (ORCA-R2, Hamamtsu Photonics, Bridgewater, NJ, USA). The light source for fluorescence imaging was provided by the X-Cite series 120 PC unit (Excelitas Technologies, Waltham, MA, USA). The microscope was surrounded by a stage incubator (Precision Plastics Inc., Beltsville, MD, USA) that maintained a temperature of 37 ^°^C and a humidity of about 60%. The device was placed on an automated X-Y microscope stage (MS-2000, Applied Scientific Instrumentation, Eugene, OR), with a secondary housing box that provided 5% CO_2_ gas with 100% humidity. All videos were taken in brightfield and GFP mode at 15.67 fps, and all long-term timelapse images were taken every 5 min for 24 h using CellSens software (Olympus America, Center Valley, PA, USA). In each experiment, bright field and fluorescence images were taken at selected positions at each time point. Time zero (t = 0) was defined as approximately one hour after sample seeding.

#### Spheroid segmentation

U-Net, a convolutional neural network architecture specifically designed for biomedical image segmentation developed in University of Freiburg ^63^, was used to segment the outlines of the tumor spheroids in videos obtained from brightfield microscopy. The model was trained with at least 25 images for each spheroid type. The spheroid area obtained from the U-Net segmentation was used to calculate the strain.

#### PAA indentation measurement

To quantify the PAA gel deformation during tumor spheroid compression, the defocused ring method was used to track the movement of the beads located at the PAA gel and tumor spheroid interface^64^. Briefly, the diameter of the defocused green fluorescent bead ring formed by PAA gel indentation during tumor spheroid compression was measured in each video frame using the Hough Circle Transformation plugin in ImageJ. The radii of the rings were then used to quantify the extent of indentation over time. This ring radius correlates directly with the depth of indentation into the PAA gel. The calibration plot of the defocused ring is shown in Fig. S8, and its measurement accuracy is validated in Fig. S9.

#### Vorticity and speed within a spheroid

The vorticity and speed within a spheroid were calculated from brightfield image sequences using the MATLAB-based PIVLab toolbox. Brightfield time-lapse images of compressed and uncompressed spheroids were imported into PIVLab, where a particle image velocimetry (PIV) algorithm was applied to compute velocity vector fields between consecutive frames. A single-pass FFT window deformation algorithm was used with an interrogation window size of 64 pixels and a step size of 16 pixels. The resulting 2D velocity fields were then used to compute the vorticity at each point, enabling visualization and quantification of rotational flow patterns and localized tissue motion in response to mechanical loading.

### Determining Young’s modulus of PDMS and PAA gel

The Young’s modulus of the PDMS used in the microrheometer was measured using a dynamic mechanical analyzer (TA Instruments DMA Q800). The results confirmed that the material exhibited linear elastic behavior, with a Young’s modulus of 1.33 ± 0.011 MPa (n = 3) (Fig. S5). The Young’s modulus of the PAA gel was measured to be 638.68 ± 41.5 Pa (n=18) using the modified indentation method developed in our labs previously ^65^, which accounts for modulus overestimation caused by the thin substrate. Briefly, a steel sphere (600 µm diameter, 7.8 g/mL density) was gently placed on the surface of the PAA gel submerged in PBS (Fig. S6). Both the top and bottom surfaces of the gel were coated with 1 µm green fluorescent beads to allow accurate measurement of gel thickness and indentation depth. The indentation depth was determined by visualizing the vertical displacement of the green fluorescent beads using a microscope. The difference in the z-position of the microscope stage with and without the metal sphere was recorded and corrected for optical distortion due to refractive index mismatch at the air-gel interface by applying a correction factor of 1.31. This correction factor was obtained experimentally in our lab previously ^65^. The indentation depth was then used to calculate the Young’s modulus using the modified Hertzian contact theory. To validate the accuracy of the indentation-based modulus measurements within our device setting, we also performed independent shear rheology measurements using a commercial rheometer (TA Instruments DHR3). The results confirmed that the PAA gel exhibits linear elastic behavior within the relevant strain range and yielded a Young’s modulus of 730.7 ± 15.9 Pa (n = 3), which closely agrees with the indentation-based value (Fig. S16). For all analyses, we used the Young’s modulus measured via the indentation method, as it was obtained under the same experimental conditions and gel layer configuration used in our microrheometer system.

### COMSOL FEM modelling

A COMSOL finite element simulation was conducted during the design phase of the microrheometer development to optimize device dimensions and material stiffness, as well as to validate the relationship between applied pressure and piston displacement (See Fig. 1D,E,F). The device geometry chosen is shown in Fig. 1D. To reduce computational load, the simulation was performed using an axisymmetric model in COMSOL. PDMS was modeled as a linear elastic material with modulus of 1.33 MPa and Poisson’s ratio of 0.49, while the tumor spheroid and PAA gels were modeled as incompressible Neo-Hookean hyperelastic materials. We note that the simulation result shown in Fig. 1E is insensitive to either the spheroid or PAA modulus, as both are significantly softer than PDMS. In the simulation, the bottom glass was modeled as a rigid material with fixed constraints, the sides of the device were fixed, and the pressure was applied to the top surface (Fig. 1E). An extra fine mesh was applied across the model to ensure numerical accuracy.

## Results and discussion

### Design and calibration of the microrheometer

The microrheometer (Fig. 1) is designed with two main objectives: (1) to characterize the mechanical properties of living microtissues in a physiologically realistic condition, and (2) to enable real-time imaging of single-cell dynamics during dynamic compression.The device fits onto a standard 75 x 25 x 1 mm glass slide and consists of six compression units and 6 control units with no compression (Fig. 1A, B). Each compression unit (Fig. 1C) is composed of three layers: the sample chamber layer (L1), the deformable PDMS membrane with a piston layer (L2), and the pressure control chamber layer (L3) (Fig. 1B, C). The deformable membrane layer (L2) and pressure chamber layer (L3) form a pressure control unit and is placed directly above the sample chamber layer. The pressure control unit provides pressure control for 6 interconnected pressure chambers (Fig. 1B) using a commercially available dynamic pneumatic pressure controller. There are 12 sample chambers in L1, of which 6 of them are control chambers, to house spheroid embedded ECM. Each sample chamber has an inner diameter of 3 mm, an outer diameter of 5 mm, and a height of H which can be varied. Four equally spaced slits of 0.5 mm width in the sample chamber wall are introduced to facilitate nutrient transport and pressure release during compression. A rectangular wall with a width of 2.5 mm and a height of H is patterned along the periphery in L1 to contain cell growth medium in the device. The 4 access ports of the medium reservoir are shown in pink in Fig. 1A.

A key component of the microrheometer is the thin polyacrylamide (PAA) gel membrane at the bottom of the cell chamber for measuring the compression force onto the spheroid directly in a physiologically realistic setting. PAA gel was selected for its well-characterized linear elastic behavior within the strain range of interest (0 - 30 %) ^66-70^. The tumor spheroid is placed on top of the PAA gel. When the spheroid is compressed, the deformation of the PAA gel is measured to calculate the compression force exerted onto the spheroid.

To calibrate the microrheometer, we applied various pressures in the pressure control unit, and measured the piston movement, Δh, and validated this result against the COMSOL calculation. The geometry of the device and the setup is shown in Fig. 1D. Experimentally, the piston movement Δh was measured by tracking the positions of the 1 µm fluorescence beads coated at the bottom of the piston with an epifluorescence microscope. In computation, a COMSOL FEM simulation was used (Fig. 1E). As shown, the experimental results closely aligned with the computation results (Fig. 1F). The piston displacement exhibited an approximately linear relationship with the applied pressure from 0 to 14 kPa (Fig. 1F). The displacement of the piston (Δh) remained nearly identical regardless of the presence or absence of a tumor spheroid, indicating that the force generated by the applied pressure is primarily absorbed by the PDMS membrane rather than the tumor spheroid.

### Strain measurement

Tumor spheroids were placed in the sample chamber and immersed in culture medium. Strain was inferred from images taken at the vertical midplane of the spheroid.

Upon compression, the spheroid underwent a cross-sectional area change as shown in Fig. 2A1,2. The cross-sectional area was segmented using a machine learning segmentation algorithm (U-Net), with yellow and red outlines representing the spheroid boundaries before and after compression, respectively. From the segmented area A, the effective spheroid radius, R, was calculated using 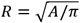. The strain is defined as *ε* = ΔR/Ro, where ΔR= R-Ro represents the change in effective radius due to compression, and Ro is the original radius before compression.

**Fig. 2.**
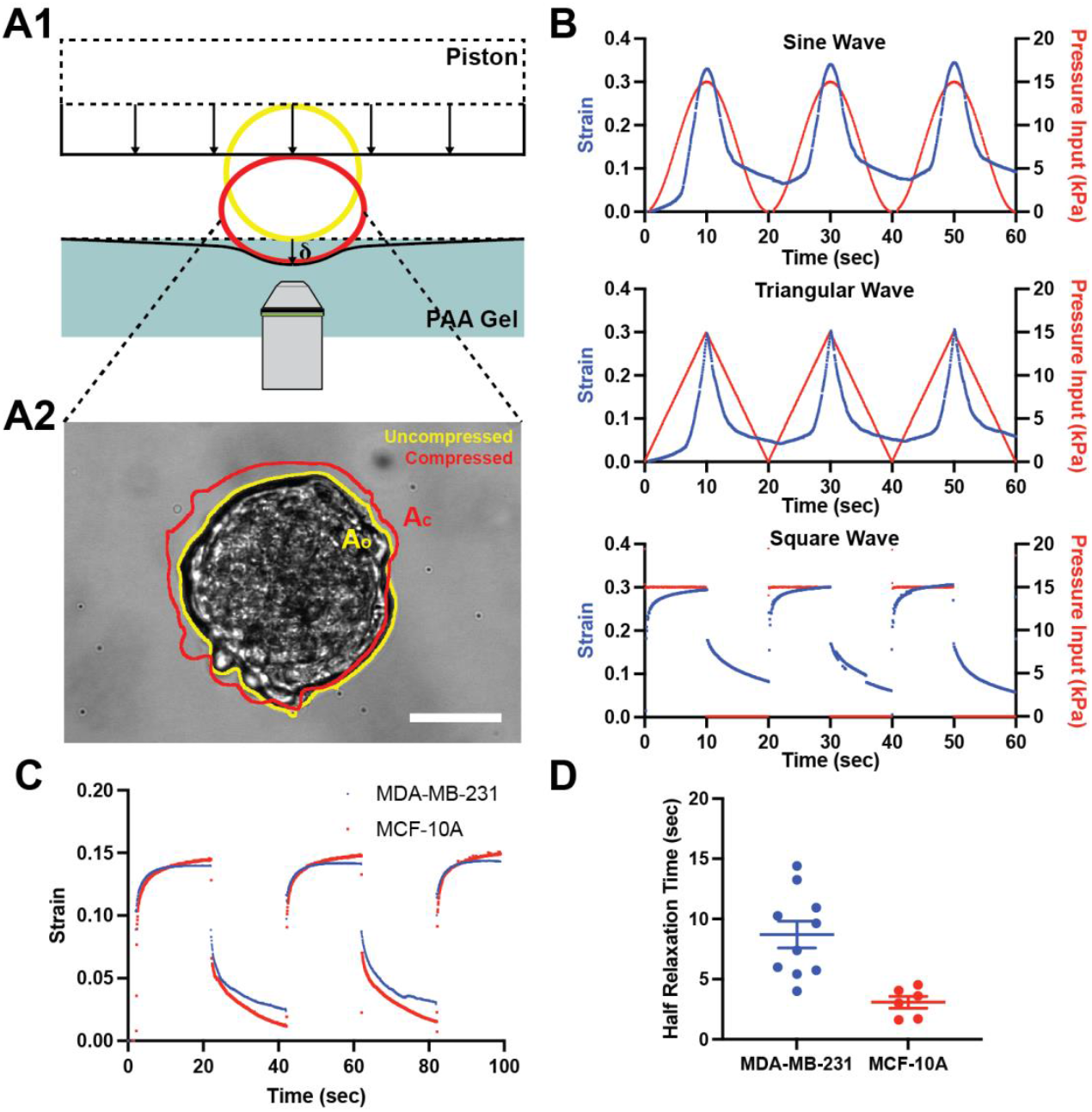
Strain measurement and the strain relaxation time of the spheroids. A1. Schematic of a tumor spheroid before (yellow, dashed lines) and during (red, solid lines) compression. The piston moves down vertically compressing a tumor spheroid underneath, resulting in the change in cross-sectional radius of the tumor spheroid. A2. A micrograph of an uncompressed MCF-10A tumor spheroid imaged at vertical midplane. The yellow outline indicates the area of the uncompressed spheroid Ao, and the redline indicates the area of the compressed spheroid, Ac. The scale bar is 100 µm. B. Spheroid strain response to sinusoidal, square, and triangular pressure wave compression with a period of 20 seconds from the microrheometer. The blue dots show the strain of the tumor spheroid obtained from bright field images with respect to time and the red dots represent the pressure applied to the pressure controller. The sampling rate is 15.67 Hz. C. Strain response of MDA-MB-231 and MCF-10A tumor spheroids when subjected to square wave compression. The maximum pressure here is 10 kPa. D. Half-relaxation time of the MDA-MB-231 and MCF-10A spheroids. Half relaxation time was defined as the time it takes for the strain to decrease to 50% of its original value after the pressure is released.

The pressure applied to the system can be controlled precisely in any function forms by the pressure controller. Sinusoidal, triangular, or square pressure waves can be transmitted to the system, resulting in strain responses as shown in Fig. 2B and SMovie 3-6. The viscoelastic properties of the MDA-MB-231 and MCF-10A tumor spheroids (Fig. 2C) can be inferred using the strain response curve of the spheroid when subjected to the square wave pressure. Here, as the compression is removed at t = 20 sec, the tumor spheroid area gradually returns to its original state. Fig. 2C compares the relaxation curves of malignant (MDA-MB-231) and non-malignant (MCF-10A) spheroids under the same strain (∼15%). It shows that the non-tumorigenic MCF-10A spheroids exhibited a faster rebound upon pressure release compared to the malignant MDA-MB-231 tumor spheroids, indicating a greater elasticity and lower viscosity in the MCF-10A spheroids (Fig. 2C). The measured half relaxation time of MCF-10A spheroids was significantly shorter than that of MDA-MB-231 spheroids (8.707 ± 1.117 sec for MDA-MB-231 spheroids and 3.086 ± 0.494 sec for MCF-10A spheroids), as shown in Fig. 2D. The half relaxation time was defined as the time required for the spheroid strain to decrease to 50% of its initial value following immediate decompression. In a linear viscoelastic model, the half relaxation time is a measure of viscosity over modulus.

Our results demonstrate that malignant tumor spheroids exhibit more fluid-like behavior compared to non-tumorigenic spheroids. The viscoelastic properties of spheroids arise from a combination of single-cell mechanical properties and intercellular adhesion. Single MCF-10A cells are known to be stiffer than MDA-MB-231 cells, and they form stronger cell-cell adhesions mediated by E-cadherin ^62^. In contrast, MDA-MB-231 cells exhibit weaker adhesion, relying primarily on indirect adhesion through the ECM ^71^. The extent to which differences in single-cell mechanics and intercellular adhesion contribute to the emergent viscoelasticity of the spheroids remains an open question for future study.

One advantage of the microrheometer is the ease with which the frequency and amplitude of the pressure wave can be precisely controlled. As shown in Fig. S7, applying a longer period pressure wave revealed a strain response curve that displayed plastic deformation over extended timescales in the tumor spheroid system.

### Stress measurement

To obtain the stress experienced by the tumor spheroid from the microrheometer, the PAA gel force sensor embedded at the bottom of the sample chambers was used. The stress is defined as the total force experienced by the spheroid divided by the cross-sectional area of the spheroid at the mid-z plane. This represents an effective or averaged stress, as the internal stress distribution within the spheroid is not uniform. When the force is applied to the tumor spheroids, the force experienced by the spheroids is transferred to the PAA gel, deforming the PAA gel by δ (Fig. 3A1). This deformation, along with the PAA gel modulus, is then used to calculate the force applied to the sample. To measure the deformation of the PAA gel, 1 µm fluorescent beads were placed at the top of the PAA gel. The defocused particle tracking method is used to measure the PAA gel deformation δ ^64^. Here, the indentation is computed using the defocused ring size change (Fig. 3A2 and Fig. S8) and is visually verified using cross sectional images shown in Fig. 3A3. The force was then calculated from the modified Hertzian Contact Theory ^65^. Briefly, the force was calculated using,

**Fig. 3.**
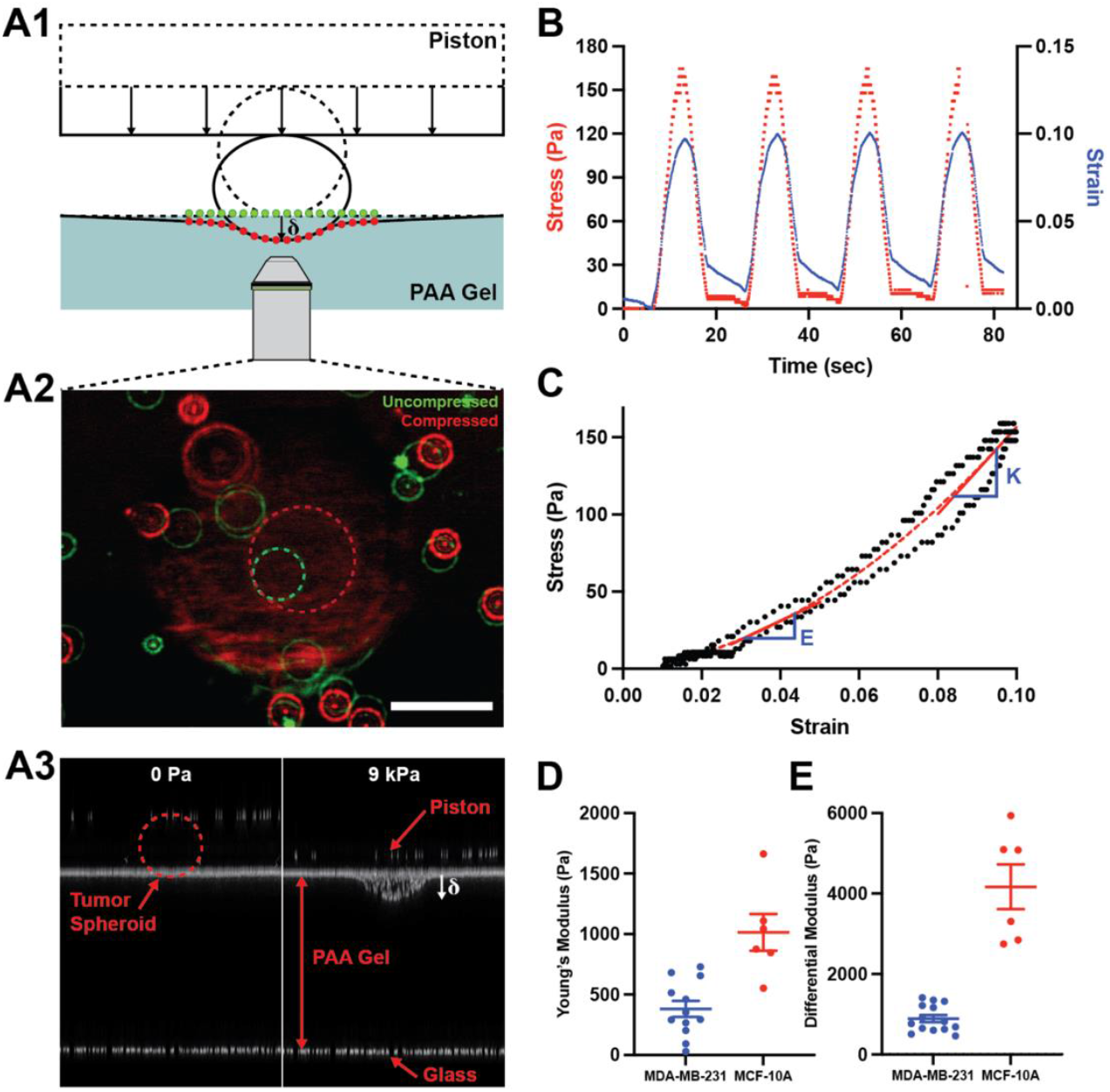
Stress measurement and the moduli of the spheroids. A1. Schematic of a tumor spheroid with and without compression. Fluorescent beads are placed on the PAA gel surface to visualize the deformation caused by the tumor spheroid. The green beads represent the bead location without compression, and the red beads represent the bead location with compression. A2. Two Epi-fluorescence micrograph images without compression (green) and with compression (red) are overlayed. The defocused rings get bigger when the beads move closer to the objective lens. This fact was used to measure the deformation of the gel. The scale bar is 100 µm. A3. A 3D projection of the z-stack images in the sample chamber with 0 Pa and 9 kPa of pressure applied to the pressure chamber. B. The stress and strain curve as a function of time for an MCF-10A spheroid. Stress is evaluated using the force measured divided by the cross-sectional area of the spheroid. Strain is ΔR/R. Four cycles of sinusoidal pressure waves with a period of 20 seconds and maximum pressure of 12 kPa were applied. C. The stress-strain response curve of an MCF-10A tumor spheroid. The slope of the curve is used to calculate the modulus of the tumor spheroid. The Young’s modulus (E) is measured in the linear region of the curve and the differential modulus (K) is measured at strain of 10%. D. The Young’s modulus (E) of MDA-MB-231 and MCF-10A tumor spheroids. E. The differential modulus (K) of MDA-MB-231 and MCF-10A tumor spheroids at 10% strain.

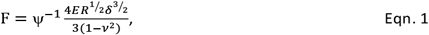

where E is the modulus of the PAA gel, R is the radius of the tumor spheroid, δ is the PAA gel deformation, and v is the Poisson’s ratio, which can be assumed as 0.5 for incompressible biological materials in short time scale. The correction factor, Ψ, was introduced by previous work from our labs to account for the thin PAA gel ^65^. Finally, the stress was acquired by dividing the force by the cross-sectional area of the tumor spheroid.

### Stress-strain curve and moduli of tumor spheroids

To characterize the mechanical properties of tumor spheroids, stress and strain were measured under sinusoidal cyclic loading with a period of 20 seconds and strain of 10% (Fig. 3B, C). The stress-strain curve in Fig. 3C revealed that (1) the tumor spheroid was nonlinear and had strain stiffening property; (2) the loading and unloading curves exhibited hysteresis, indicating the viscoelastic behavior of the tumor spheroids. Using the vertically averaged, and interpolated stress-strain curve (dotted red midline in Fig. 3C), we computed the Young’s modulus of the spheroid, the slope of the linear region of the curve near low strain. In addition to the Young’s modulus, we also calculated the differential modulus, defined as the local slope of the stress-strain curve at the 10% strain. The differential modulus provides a measure of the instantaneous stiffness of the spheroids locally around the specific strain. This is important because almost all biological materials exhibit nonlinear mechanical properties ^25^.

The measured average Young’s modulus values were 380.6 ± 66.3 Pa for MDA-MB-231 spheroids and 1014.6 ± 151.8 Pa for MCF-10A spheroids (Fig. 3D), while the differential modulus at 10% strain was 893.2 ± 89.6 Pa for MDA-MB-231 spheroids and 4168.8 ± 557.4 Pa for MCF-10A spheroids (Fig. 3E). These results are obtained using 14 spheroids for MDA-MB-231 and 6 spheroids for MCF-10A spheroids. We note that force measurements using PAA gel can be inaccurate when the stiffness of the spheroid is comparable to that of the PAA gel. This error arises from the violation of the assumptions in the Hertzian contact theory, which assume a rigid spherical indenter deforming an elastic substrate. When the spheroid is not significantly stiffer than the substrate gel, this assumption is violated, introducing error into the calculated modulus. To evaluate this effect, we performed a finite element analysis and confirmed that, within the stiffness ratio range relevant to our experiments (E_Sph._/E_PAA_ ϵ [1,3]), this error remains below 21% (see section 2 of the supplementary materials and Fig. S15). After applying the correction factor derived from this analysis, the corrected Young’s modulus values were 396.8 ± 60.1 Pa for MDA-MB-231 spheroids and 903.0 ± 111.9 Pa for MCF-10A spheroids. Future improvements can be made through: (i) the use of a softer PAA gel relative to the spheroid sample; (ii) experimental validation of the correction factor using a soft spheroid with known mechanical properties.

Here, we show for the first time that tumor spheroids are both strain-stiffening under compression regardless of its malignancy state. This differs from behavior observed in collagen matrices, where collagen fibers buckle (or soften) under compression. This strain stiffening thus is likely contributed by the cell mechanics. Furthermore, we also show that the more malignant MDA-MB-231 spheroids are significantly softer than the non-malignant MCF-10A spheroids, consistent with previous modulus measurements of single cells and spheroids obtained using atomic force microscopy and microtweezers ^28, 72^. This, however, differs from the clinical observations that the tumors are a lot stiffer than the normal tissues. This discrepancy may arise from the fact that clinical tumor stiffness reflects not only the mechanical properties of cancer cells, but also contributions from the surrounding extracellular matrices which often undergo collagen crosslinking and fibrosis in malignant state ^6, 73, 74^. Therefore, while individual tumor cells or spheroids may be softer, the surrounding tumor stroma can dominate the bulk tissue mechanics, leading to the elevated stiffness detected in vivo. Further mechanical testing using spheroid embedded ECM can answer this important question.

### Tumor cell dynamics under compression

The transparent design of the microrheometer enables live-cell imaging, making it an ideal tool for correlating tumor mechanics with tumor cell dynamics. Here, we studied tumor cell dynamics both within the spheroids and outside the spheroids under compression. Note that we used a device with the dimensions shown in Fig. 1D, except with the sample chamber height (H) set to 500 µm and without the PAA gel layer (See Fig. S10). The tumor spheroids embedded in ECM were loaded directly at the bottom of the sample chamber (top 6 wells with MDA-MB-231 spheroids, and bottom 6 wells with MCF-10A spheroids), and the collagen was allowed to polymerize for 45 minutes in 37^°^C incubator before experiment. We note that the same DMEM/F12 medium is used for all the experiments presented here.

#### Compression moderately modulates tumor spheroid protrusions into ECM

To investigate the effects of compression on tumor spheroid invasion into the ECM, we followed tumor cell dynamics under compression. Tumor spheroids in the experimental sample chamber (left six wells) were subjected to 30% Δh/h compression, while those in the control sample chambers (right six wells) remained uncompressed for 24 hours.

Under these conditions, distinct morphological responses were observed: malignant MDA-MB-231 spheroids exhibited sharp protrusions into the ECM, while non-tumorigenic MCF-10A spheroids stayed mostly circular (Fig. 4). Full time-lapse movies are available in SMovie 7-10. Consistent with previous studies ^75, 76^, the malignant (MDA-MB-231) spheroids showed significantly greater invasion into the ECM than non-tumorigenic (MCF-10A) spheroids. Using images shown in Fig. 4A1,2 and Fig. 4B1,2, we computed spheroid area fold increase as well as circularity. Fig. 4A3, B3 showed that compression moderately reduced the spheroid area fold change over time for both the MDA-MB-231 and the MCF-10A spheroids within the 24-hour observation window. The circularity, defined as *c*= 4*π (Area/Perimeter*^*2*^),was also calculated from the outlines of each tumor spheroid. The circularity of the MDA-MB-231 spheroids dropped significantly faster than that of the MCF-10A spheroids as the malignant MDA-MB-231 cells protruded into the ECMs. For the malignant MDA-MB-231 spheroids, there was no significant difference in circularity between the compressed and the uncompressed groups (Fig. 4A4) as both groups exhibited substantial protrusions. For the MCF-10A spheroids, the circularity was lower for the uncompressed group than the compressed group, which indicated that invasion was reduced due to compression for MCF-10A spheroids (Fig. 4B4), consistent with the message from Fig. 4B3.

**Fig. 4.**
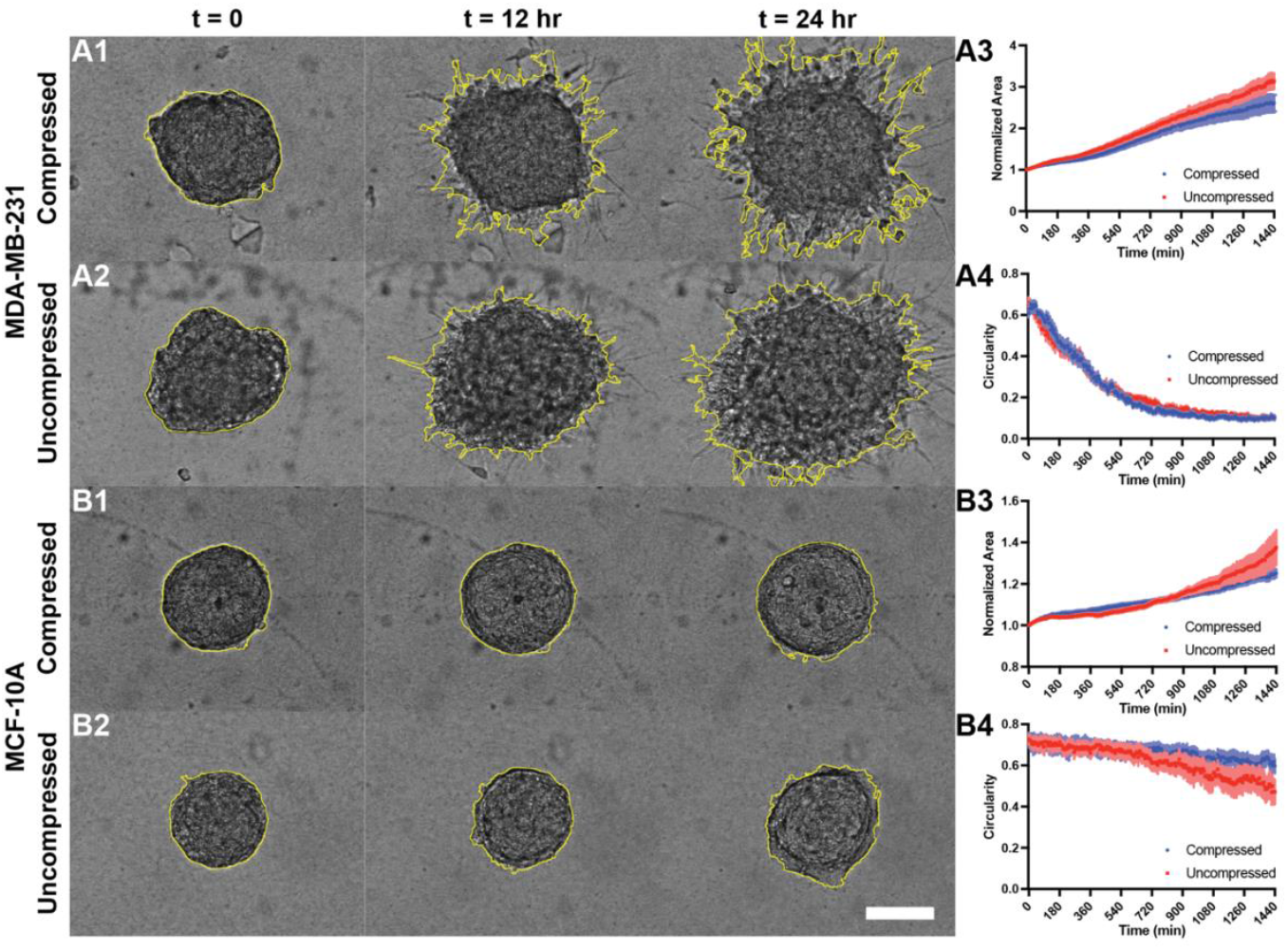
Compression moderately reduces tumor spheroid area expansions over time. A1-2 (B1-2). Time-lapse brightfield images of MDA-MB-231 (MCF-10A) tumor spheroids embedded in 3.5 mg/mL collagen under compressed and uncompressed conditions. The scale bar is 100 µm. A3 (B3). Normalized spheroid coverage area of MDA-MB-231 (MCF-10A) spheroids under compressed and uncompressed conditions over 24 hours. A4 (B4). Circularity of MDA-MB-231 (MCF-10A) spheroids for compressed and uncompressed conditions. The darker dots represent mean values, and the error bars indicate SEM at each time point. These values were calculated from 11 compressed and 10 uncompressed MDA-MB-231 spheroids, and 12 compressed and 8 uncompressed MCF-10A spheroids.

Results in Fig. 4 show that MDA-MB-231 spheroids invade significantly more into the ECM than the non-tumorigenic MCF-10A spheroids. This is consistent with the fact that malignant MDA-MB-231 cells, but not MCF-10A cells, have a high level of integrins, needed to invade into the ECMs. In contrast, compression moderately reduced the invasion of both types of spheroids into ECM. We conjecture that compression here leads to compaction of collagen matrices surrounding the spheroids, which may pose a mechanical barrier to cell invasion. Future imaging of the ECM architecture will be required to verify this conjecture.

#### Compression induces vortex motion within non-tumorigenic tumor spheroids

To understand the roles of compression on the tumor spheroid fluidity, we followed single cell dynamics within the spheroids when they were subjected to the 30% Δh/h compression. Interestingly, we discovered that non-tumorigenic tumor cells form a vortex within the spheroid under compression, while this vortex was absent in malignant tumor spheroids. To characterize the internal cellular motion within the tumor spheroids, particle image velocimetry (PIV) was performed on time-lapse brightfield images using MATLAB (PIVLab) (Fig. 5A1-2, B1-2). The full time-lapse velocity vector fields from PIV analysis are available in SMovie 11 and 12.

**Fig. 5.**
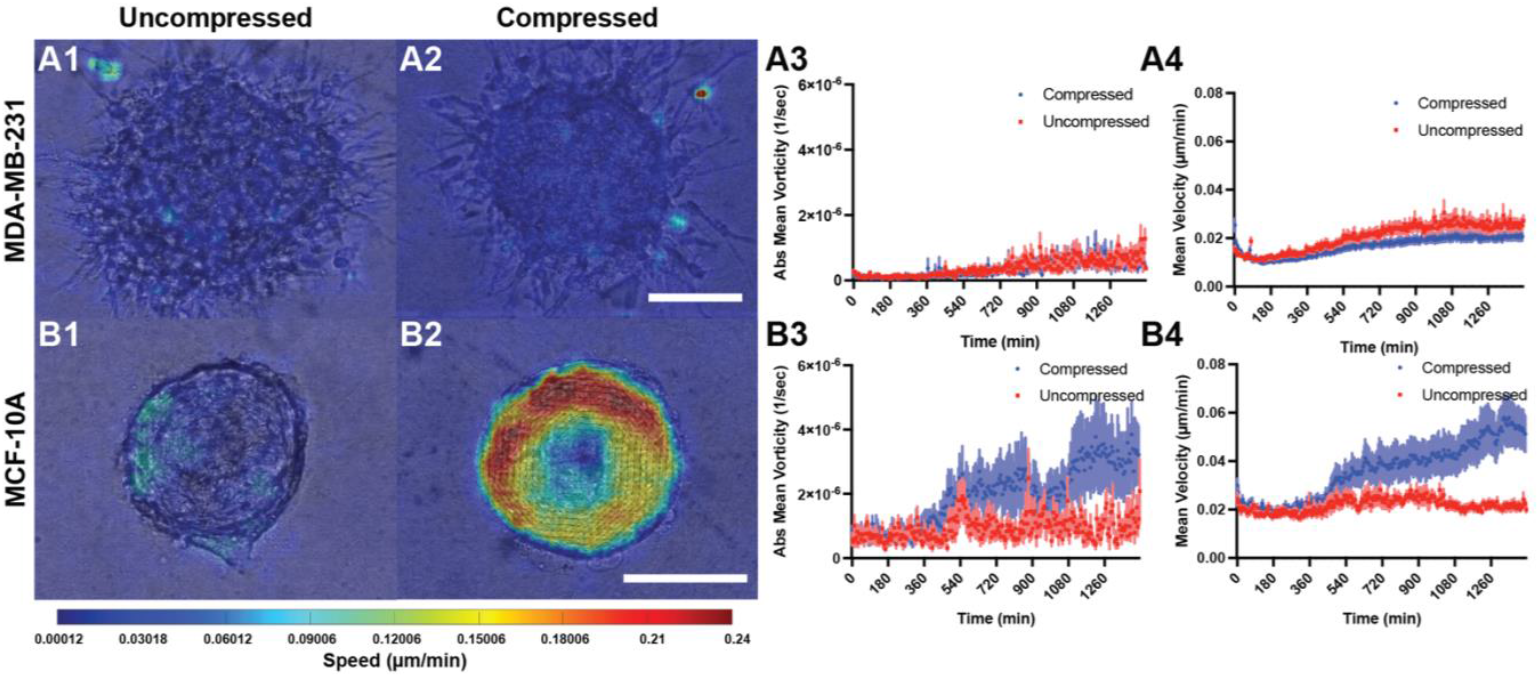
Compression induces internal cellular vortex motion within non-malignant tumor spheroids, but not in malignant tumor spheroids. A1-A2. Velocity vectors of MDA-MB-231 cells under uncompressed (A1) and compressed (A2) conditions. A3. Absolute mean vorticity of the MDA-MB-231 spheroids at each time point for the compressed and uncompressed conditions. A4. Mean speed of the MDA-MB-231 cells at each time point for the compressed and uncompressed conditions. B1-B2. Velocity vectors for MCF-10A cells under uncompressed (B1) and compressed (B2) conditions. B3. Absolute mean vorticity of the MCF-10A spheroids at each time point for the compressed and uncompressed conditions. B4. Mean speed of the MCF-10A tumor cells at each time point for the compressed and uncompressed conditions. The velocity field is obtained from brightfield time-lapse images using a particle image velocimetry (PIV) analysis algorithm in MATLAB. Scale bars are 100 µm. The vorticity and speed were calculated from velocity vector fields. The darker dots represent the mean values, and the error bars indicate SEM at each time point. The average lines and SEM error bars are calculated from 12 compressed and 10 uncompressed MDA-MB-231 spheroids and 14 compressed and 8 uncompressed MCF-10A spheroids.

The velocity vector fields revealed that rotational motion emerged exclusively in compressed MCF-10A spheroids, with coherent spiral patterns appearing within the first 8 hours of compression. The peripheral cells moved tangentially around the spheroid core while central cells remained stationary (Fig. 5B2), suggesting a vortex-like flow with cell motion increasing radially, a characteristic commonly observed in fluid dynamics. In contrast, no rotational motion was observed in uncompressed MCF-10A spheroids or in any MDA-MB-231 spheroids, regardless of compression.

The mean speed and mean vorticity of malignant and non-tumorigenic cells responded differently to compression, consistent with observations from the velocity field. Here, the speed is defined as 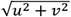, and vorticity as 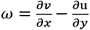,where u and v are x- and y-components of the velocity, respectively (Fig. 5A3-4, B3-4). Because the rotational direction varied between spheroids, the magnitude of vorticity was used for comparison. The mean speed and vorticity were averaged using all the velocity vectors obtained at one time point. The mean vorticity results shown in Fig. 5A3 and B3 support the visual observations of the velocity field, confirming the emergence of a coherent vortex in non-tumorigenic spheroids (MCF-10A) approximately 8 hours after compression. The mean speed of MDA-MB-231 cells decreased under compression (Fig. 5A4), whereas the mean speed of MCF-10A cells increased significantly under compression (Fig. 5B4). In compressed MCF-10A spheroids, the development of a coordinated unidirectional rotational flow promoted faster internal cellular movement. In contrast, there was a slight increase in speed observed in uncompressed MDA-MB-231 spheroids.

The emergence of vortex motion within non-tumorigenic MCF-10A spheroids under compression is consistent with the reports on unjamming transitions in epithelial layers ^77^. It has been shown that confluent epithelial layers become more fluid-like under compression. We know that MCF-10A cells have high level of cell-cell adhesion molecules, E-cadherin, which links neighboring cells together. In this case, strong intercellular adhesion enables the coordination necessary for large-scale collective motion, allowing cells to reorganize under external stress. By contrast, MDA-MB-231 cells express low levels of E-cadherin ^78-81^, leading to insufficient mechanical coupling between neighboring cells and thus lack the coordinated motion obtained in MCF-10A spheroids (Fig. 5A3). The emergence of a coherent vortex-like motion further implies a fluidization of the spheroid structure, where cells can rearrange relative to one another. Recent theoretical work has shown that fluidized spheroids are more likely to remodel the surrounding ECM architecture ^82^, such that the ECM aligns along the circumferential direction, inhibiting tumor cell invasion. Our results also indicate that non-tumorigenic spheroids have better adaptability to their surrounding environment than malignant tumor spheroids, which may prove to be important for the homeostasis of healthy tissue. Further work is underway to illustrate the roles of fluidity in tumor physiology.

## Conclusions and future perspectives

A central contribution of this work is the development of a microrheometer capable of simultaneously characterizing the mechanical properties of tumor spheroids and optically following single-cell dynamics in a physiologically realistic setting. The key innovation of the device lies in the integration of a polyacrylamide gel-based force sensor, enabling direct measurement of tissue mechanical properties. By varying the dimensions of the device, the microrheometer can be easily extended to study nuclear and single cell mechanics, and a broad range of other cell/tissue types.

Using this device, we found that non-tumorigenic (MCF-10A) and malignant breast tumor (MDA-MB-231) spheroids are both nonlinear and viscoelastic, exhibiting a strain stiffening behavior. The measured Young’s modulus of MCF-10A and MDA-MB-231 spheroids was consistent with previously reported values. By enabling simultaneous mechanical measurement and live imaging, the microrheometer revealed that compression can induce large-scale motion in non-tumorigenic spheroids, but not in malignant spheroids. In addition, compression moderately reduced invasiveness when tumor spheroids were embedded in 3.5 mg/mL collagen.

Looking ahead, the microrheometer can find wide applications in both applied and basic research. Its ability to measure tissue stiffness within minutes opens doors for clinical applications including biopsy specimens and patient-derived organoids. It can be used as a standard tool for mechanical phenotyping in personalized medicine. On the basic research front, the microrheometer can be integrated with molecular and biochemical readouts. Introducing fluorescent markers of mechanotransduction pathways (e.g., YAP/TAZ, actin/myosin), it would enable direct correlation between mechanical loading and molecular responses. RNA sequencing analysis from compressed versus uncompressed spheroids could further reveal mechanical stress-induced gene expression changes. In parallel, exploring dynamic loading – such as sinusoidal or cyclic compression – could better mimic in vivo conditions like fluctuating mechanical forces due to breathing, muscle contractions, blood pressure pulsations, and organ motion ^83^. These studies may uncover how tumors adapt to fluctuating mechanical environments and reveal new regulatory mechanisms.

## Author contributions

YJS and MW created and designed the research project. YJS carried out the device fabrication and experiments. YJS and ML performed data analysis. YJS, BZ, JK, NB, and CYH carried out the FEM simulation. BZ and CYH carried out the theoretical calculation shown in the supplementary information. CR and YJS carried out the PIV analysis. The manuscript was collaboratively written by YJS, JES, and MW, with contributions from all authors.

## Data availability

Data sets generated during the study are available from the corresponding author on reasonable request.

## Acknowledgements

This work was supported by a grant from the National Institute of Health (Grant No. R01CA221346). The authors thank Dong Wang for helpful discussions on PAA gel synthesis and Yukun Sun for helpful discussions on PIV analysis. The authors also thank Tom Pennell, Chris Alpha, Jeremy Clark, Michael Skvarla, Aaron Windsor, and Garry Bordonaro from CNF for their help in the fabrication steps. This work was performed in part at the Cornell NanoScale Facility, a member of the National Nanotechnology Coordinated Infrastructure (NNCI), which is supported by the National Science Foundation (Grant NNCI-2025233). JES is the Betty and Sheldon Feinberg Senior Faculty Scholar in Cancer Research.

