## Supplementary Information for "A microfluidic rheometer for tumor mechanics and invasion studies"

### 1. Supplementary Figures

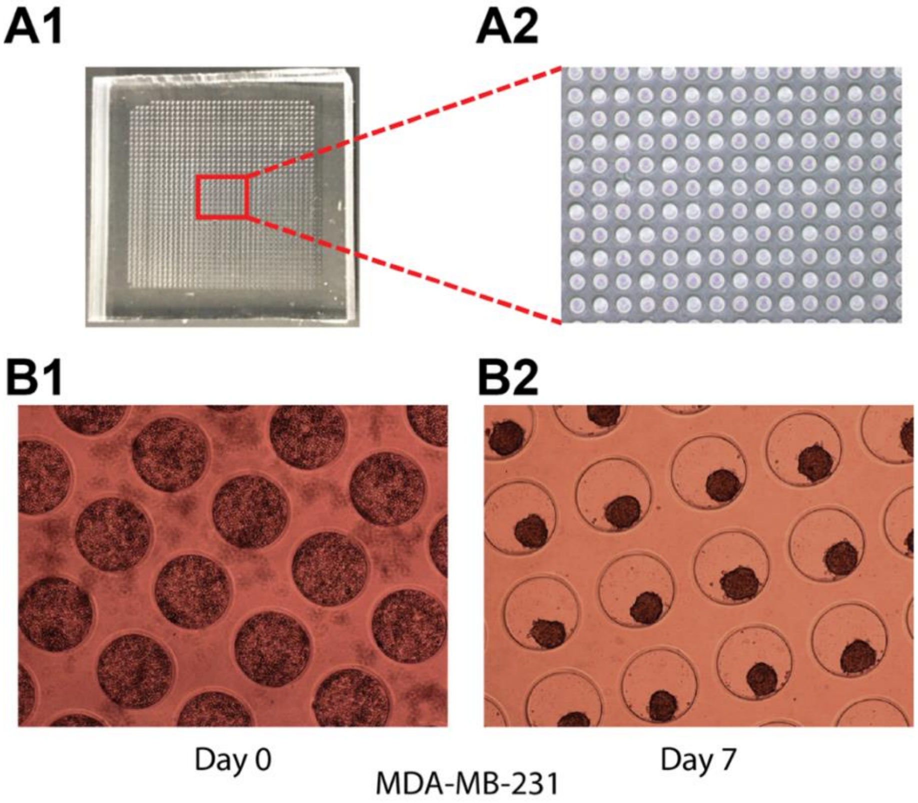


**Figure S1: Microwell array for making tumor spheroids. A1.** Photograph of the microwell array device containing a 36 x 36 grid of microwells, each 200 µm in diameter and 200 µm in depth. **A2.** Brightfield micrograph showing a magnified view of the microwells. **B1.** MDA-MB-231 cells seeded in the microwells on Day 1. **B2.** Formation of uniform spheroids by Day 7 within the microwells.

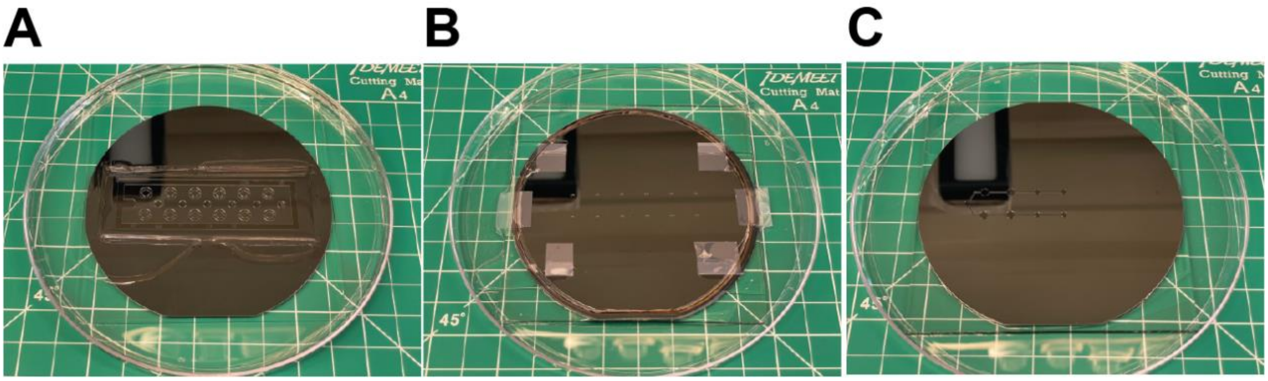


**Figure S2: Silicon masters for the microrheometer. A.** Silicon master for the sample chamber layer (L1). **B.** Silicon master for the deformable piston membrane layer (L2). **C.** Silicon master wafer for the pressure control chamber layer (L3). All the silicon masters are 100 mm in diameter.

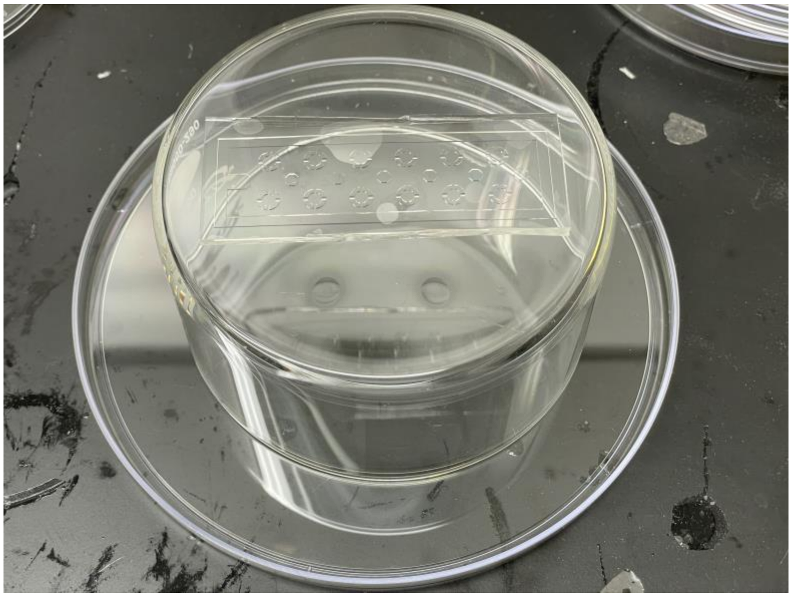

**Figure S3: In-house FOTS coating chamber.** The sample to be coated with FOTS was affixed to the bottom of a glass petri dish using double-sided tape. Four droplets of FOTS were placed on the bottom of a 150 mm plastic petri dish. The glass petri dish was then inverted and positioned over the plastic petri dish so that the sample was suspended 6 cm above the FOTS droplets, allowing vapor-phase deposition.

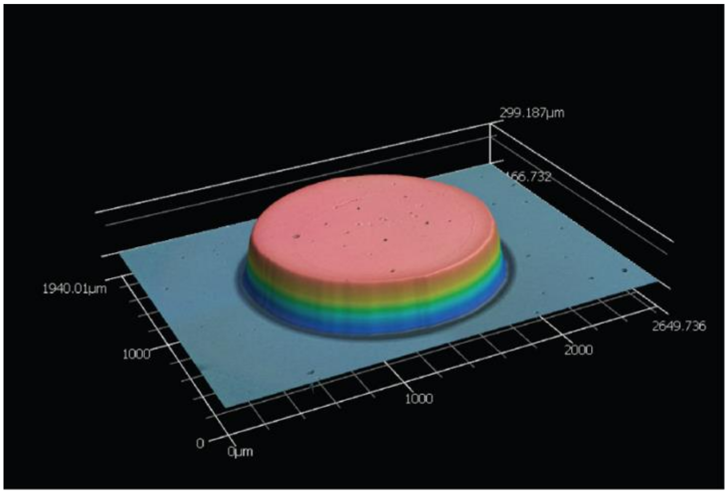


**Figure S4: Optical profilometry measurement of PDMS piston height.** A 3D laser scanning profilometer image of the PDMS piston is taken using the Keyence VK-X260 Laser-Scanning Profilometer. This confirms the piston height is 300 µm.

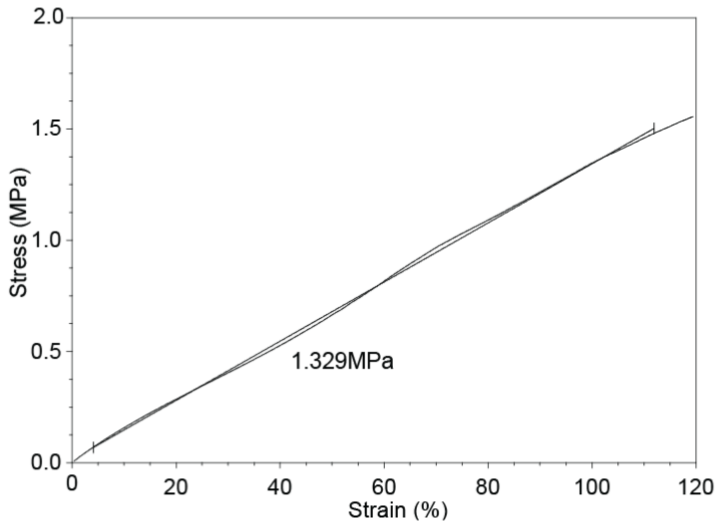


**Figure S5: Stress-strain curve of PDMS.** Stress-strain curve of a 10:1 polydimethylsiloxane (PDMS) sample under uniaxial tensile loading. The slope of the linear region indicates a Young's modulus of approximately 1.33 MPa.

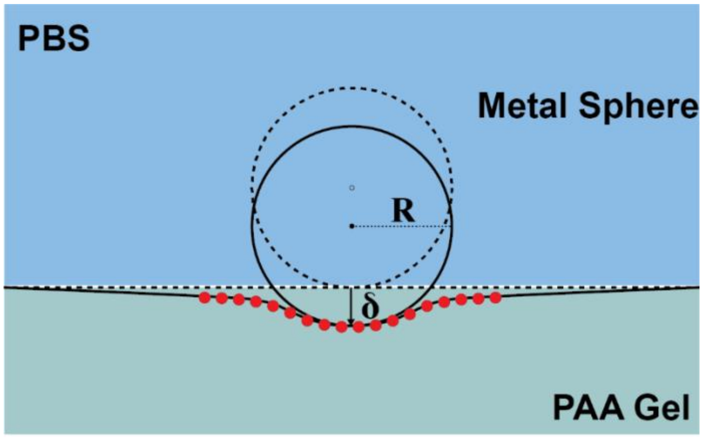


**Figure S6: Schematic of indentation-based stiffness measurement of the PAA gel using a metal sphere.** The Young's modulus of the PAA gel was measured using an indentation method previously developed in our lab [262]. A metal sphere of known radius R (600 µm diameter, 7.8 g/mL density) was gently placed onto the surface of a PAA gel submerged in PBS. The sphere caused an indentation depth $\delta$, which was used to calculate the stiffness of the gel. Red dots represent the fluorescent beads on the deformed gel surface.

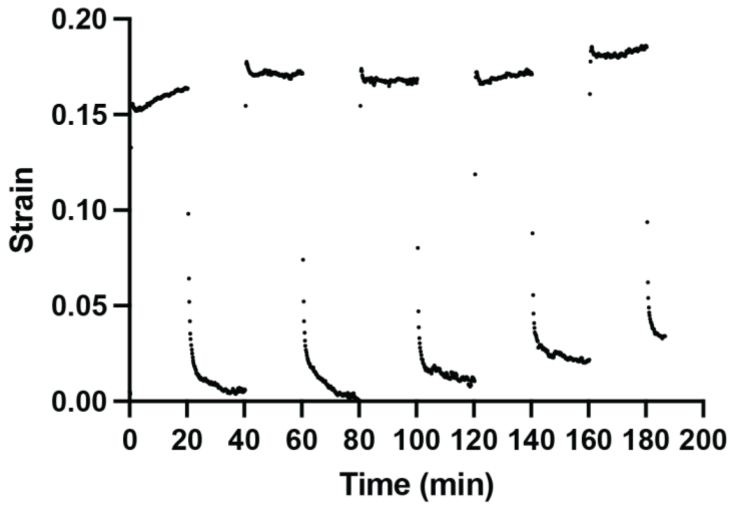


**Figure S7: Long-term compression response of MDA-MB-231 spheroids.** The radius strain of an MDA-MB-231 tumor spheroid was recorded under repeated square wave compression cycles over a 3-hour period. The period of each cycle was 40 minutes. The sampling rate is 0.1 Hz.

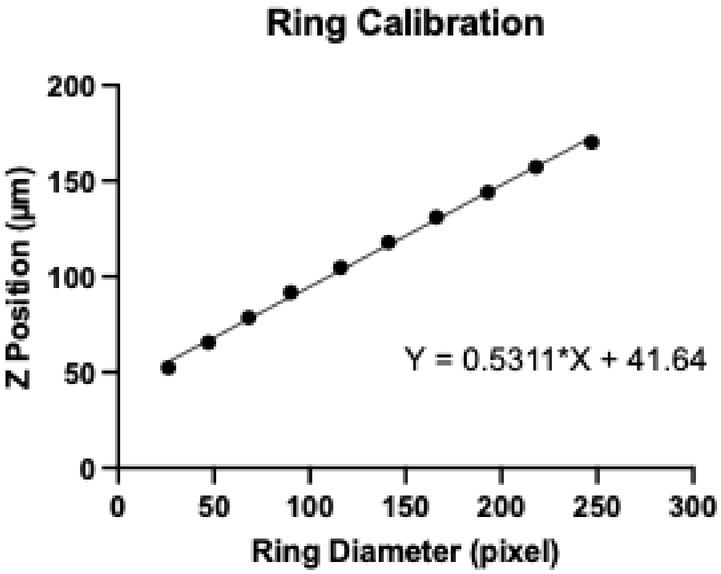


**Figure S8: Defocused ring calibration.** A z-stack of a stationary fluorescent bead placed on a PAA gel surface were imaged with a 20x objective with a 1 µm interval. The collar on the objective was turned to 2 to enhance the defocused ring effect.

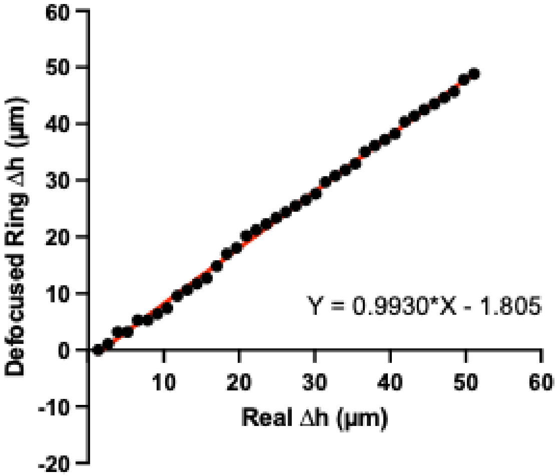


**Figure S9: Validation of displacement measurements using 3D defocused particle tracking method.** Here, the real Δh is measured using an automatic microscope stage z-axis reading and value along y axis is the displacement measured using the defocused particle method. A slope of 0.993 shows the accuracy of the defocused particle tracking method in the bead position measurement range of our experiment.

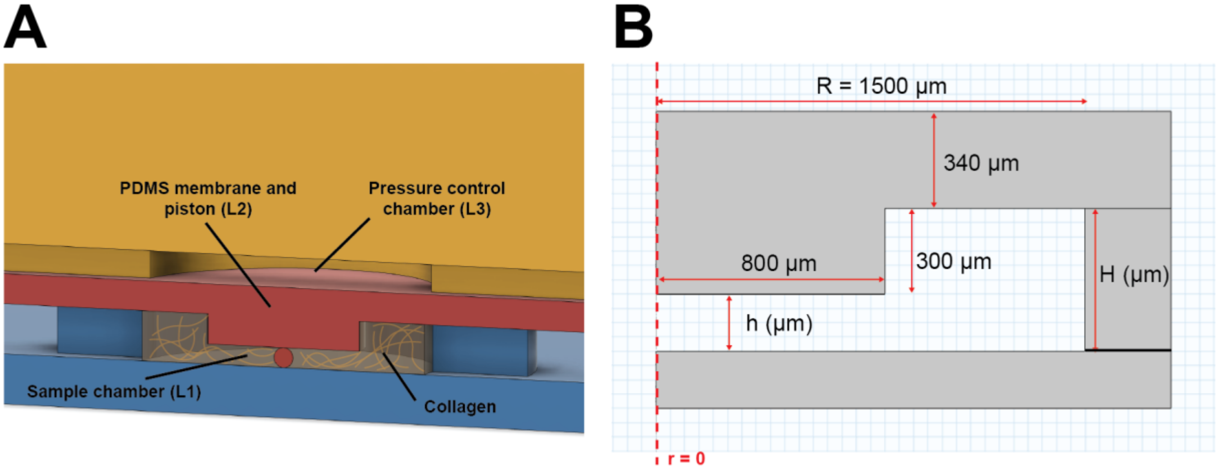


**Figure S10: Modified microrheometer for dynamic compression and invasion experiments. A.** Schematic illustration of the 3-layer microrheometer device. The bottom layer (L1) contains the modified sample chamber with a 500 μm height to accommodate the reduced height from the removal of the PAA gel. The piston membrane layer (L2) and the pressure control chamber layer (L3) remain unchanged. **B.** The dimensions of an axisymmetric compression unit. The dashed line represents the central axis of the compression unit.

### 2. Influences of spheroid modulus on the modified Hertzian contact theory calculation revealed by a Finite-Element Analysis

The modified Hertzian contact theory (Eq. 1) used to calculate the force exerted on the spheroid assumes that the spheroid modulus is significantly larger than that of the PAA gel. However, this assumption does not always hold in our experimental system. To estimate the resulting error, we performed finite element analysis (FEA) to simulate spheroid indentation across a range of spheroid stiffness values using the same gel geometry and boundary conditions as in our experiments.

We define the stress as $\sigma = F/A = F/(\pi R^2)$ , where $F$ is the applied compressive force, $R$ is the spheroid radius and $A$ is its equatorial cross-sectional area. The force $F$ is computed from contact theory as,

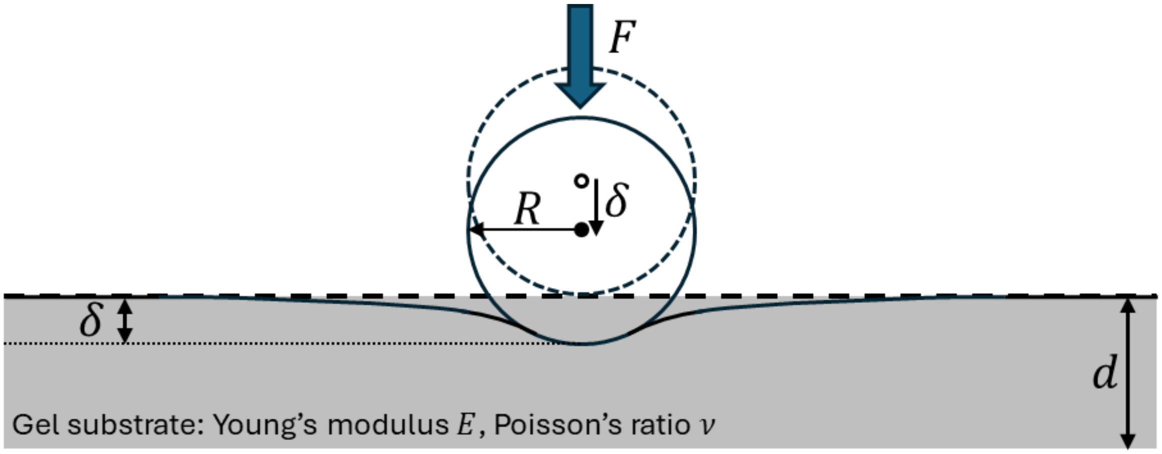


**Figure S11: Schematic illustration of Hertzian contact theory.** A rigid sphere of radius R indents a substrate of modulus E under an applied force F.

$$F = \psi^{-1} \frac{4ER^{\frac{1}{2}}\delta^{\frac{3}{2}}}{3(1-\nu^2)} \qquad \text{(Eq. S1)}$$

where $E$ and $\nu$ are the Young's modulus and Poisson's ratio of the gel substrate (we take $\nu = 0.5$ for incompressibility), $R$ is the radius of the spherical indenter, and $\delta$ is the indentation depth. The nonlinear, finite substrate thickness correction factor $\psi$ for frictionless contact, introduced by Long *et al.* [1], is

$$\psi = \frac{1 + 2.3\omega}{1 + 1.15\omega^{\frac{1}{3}} + \alpha\left(\frac{R}{d}\right)\omega + \beta\left(\frac{R}{d}\right)\omega^2} \qquad \text{(Eq. S2a)}$$

$$\omega = \left(\frac{R\delta}{d^2}\right)^{\frac{3}{2}}, \quad \alpha\left(\frac{R}{d}\right) = 10.05 - 0.63\sqrt{\frac{d}{R}}\left(3.1 + \frac{d^2}{R^2}\right), \beta\left(\frac{R}{d}\right) = 4.8 - 4.23\frac{d^2}{R^2} \qquad \text{(Eq. S2b)}$$

where $h$ is the gel thickness. Equations (Eq.2a–b) were calibrated against simulations for $0.5 \leq R/d \leq 12.7$ and $\delta/d \leq \min(0.6, R/d)$.

In our experiment, the compressing body is a soft tumor spheroid with $E_{sph.}$ rather than a rigid indenter. Under compression, the spheroid radius changes by $\Delta R$. The contact force $F$ is estimated using Eq. 1. We obtain the spheroid's Young's modulus, $E_{cal.}$ from the slope of the stress $F/A$ versus strain $\Delta R/R$ curve in the small-strain regime. Finally, we compute the ratio of $E_{sph.}$ and $E_{cal.}$.

Two considerations in this modulus estimation motivate verification with finite-element analysis. First, Equations (Eq.1–2) assume a *rigid* spherical indenter, whereas in our system the spheroid modulus can be comparable to that of the PAA gel. The rigid-indenter assumption may therefore be violated. Second, the true stress and strain fields inside the spheroid are nonuniform. Our definitions of stress $F/A$ and strain $\Delta R/R$ should be interpreted as averaged measures. Consequently, we use simulations to test whether the stress $F/A$ versus strain $\Delta R/R$ slope recovers the modulus of the tumor spheroid and to quantify any systematic bias (relative error).

To address the above concerns, we performed a series of axisymmetric 2D finite-element simulations closely matching the experimental setup.

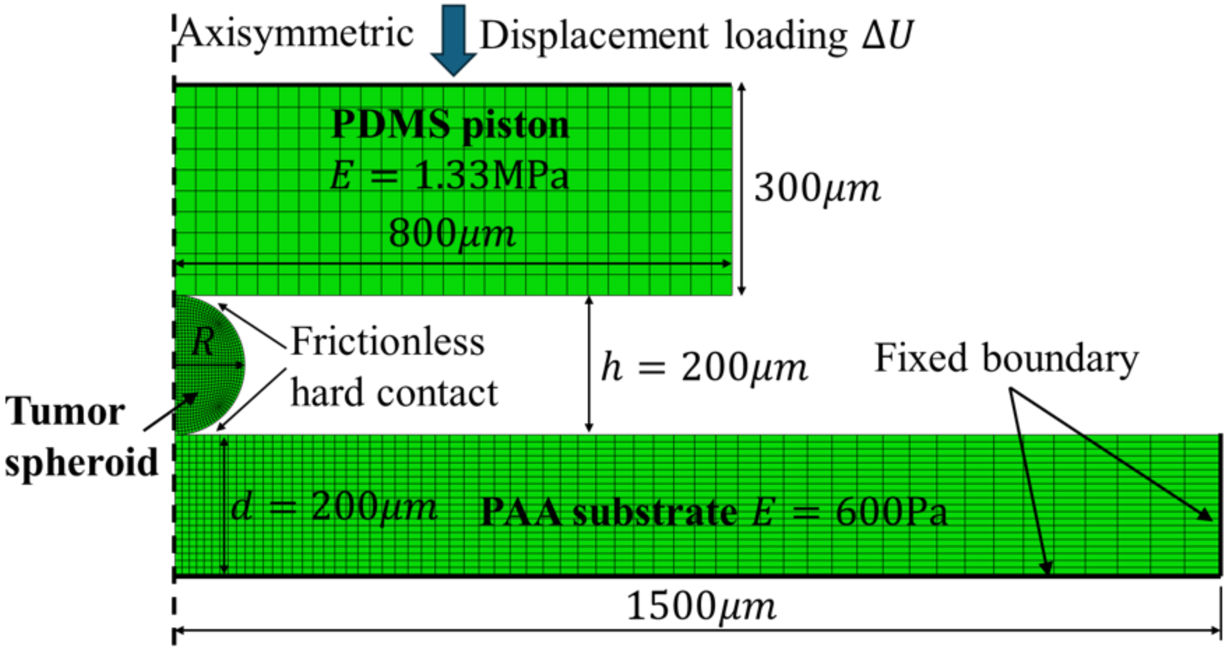


**Figure S12: Schematic of the finite-element simulation setup.** A spheroid of radius R was compressed between a PDMS piston and a PAA substrate. A vertical displacement of $\Delta U = R$ was gradually applied to the upper surface of the PDMS piston, while the bottom and lateral surfaces of the PAA substrate were fixed. The Young's moduli were taken as 1.33 MPa for PDMS and 600

Pa for PAA, and the spheroid modulus was varied. All three materials were modeled as incompressible neo-Hookean hyperelastic solids, and surface contacts were assumed frictionless.

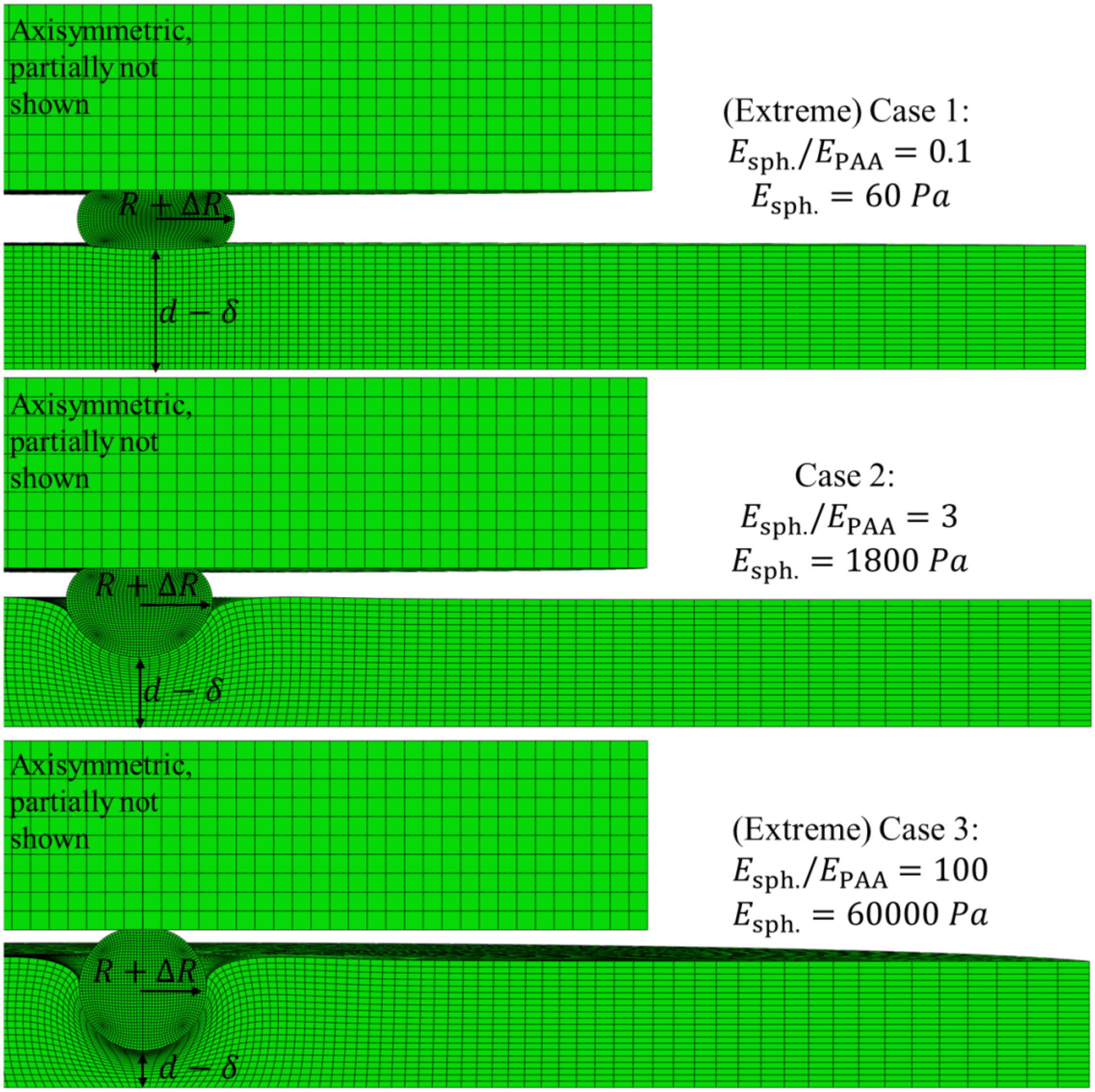


**Figure S13: Deformed shapes of the system for three different spheroid moduli from FEA computation.** Three different cases were computed using soft, intermediate and stiff spheroids in comparison to PAA gel.

The deformed configurations for three representative cases are shown in Fig. S13. Because the modulus of PDMS is much higher than that of both the spheroid and the PAA substrate, it is reasonable to treat the piston as rigid, its exact modulus has negligible influence on the overall deformation. The ratio of the spheroid modulus, $E_{sph.}$, to the PAA modulus, $E_{PAA}$, provides a dimensionless parameter that characterizes the deformation mode. When this ratio is small, the spheroid is approximately compressed between two rigid plates (Case 1 in Fig. S13). When the ratio is large, the spheroid behaves as a rigid spherical indenter, and nearly all deformation occurs in the substrate (Case 3 in Fig. S13). For the experimental conditions, the modulus ratio is on the order of unity, so both the spheroid and the PAA substrate undergo significant deformation (Case 2 in Fig. S13).

In all simulations, the indentation depth $\delta$ and the change in spheroid radius $\Delta R$ were recorded under different imposed displacements $\Delta U$. The force $F$ was then computed from Eq. 1. For each case, we plotted the stress–strain curve, defined as $F/A$ versus $\Delta R/R$, where the spheroid modulus used in the finite-element simulation is denoted $E_{\mathrm{sph}}$.

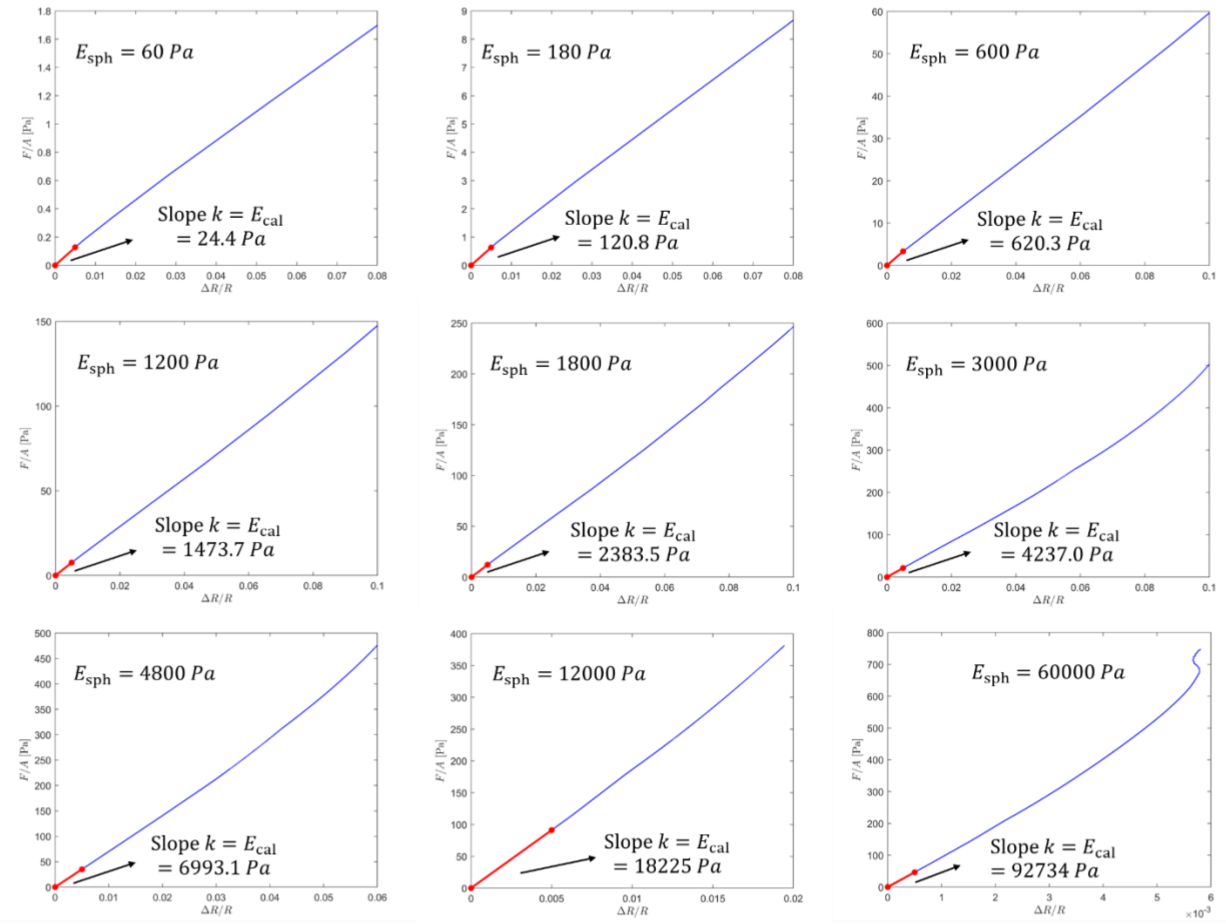


***Figure S14:*** **Stress–strain curves ($F/A$ versus $\Delta R/R$) for different spheroid moduli $E_{sph}$.** Comparison of calculated spheroid modulus $E_{cal}$ (or the slope) versus the actual spheroid modulus $E_{sph}$. the difference decreases as the spheroid modulus is significantly larger than the PAA modulus. $k$ was determined between the origin and the point where $\Delta R/R = 0.5\%$. Except for the case of $E_{sph} = 60000Pa$, in which the linear region is smaller, we determined the slope between the origin and the point where $\Delta R/R = 0.05\%$. We interpret this slope as the calculated modulus of the spheroid, $E_{cal}$. In our experiments, the modulus is likewise obtained from the slope of the $F/A$ versus $\Delta R/R$ curve.

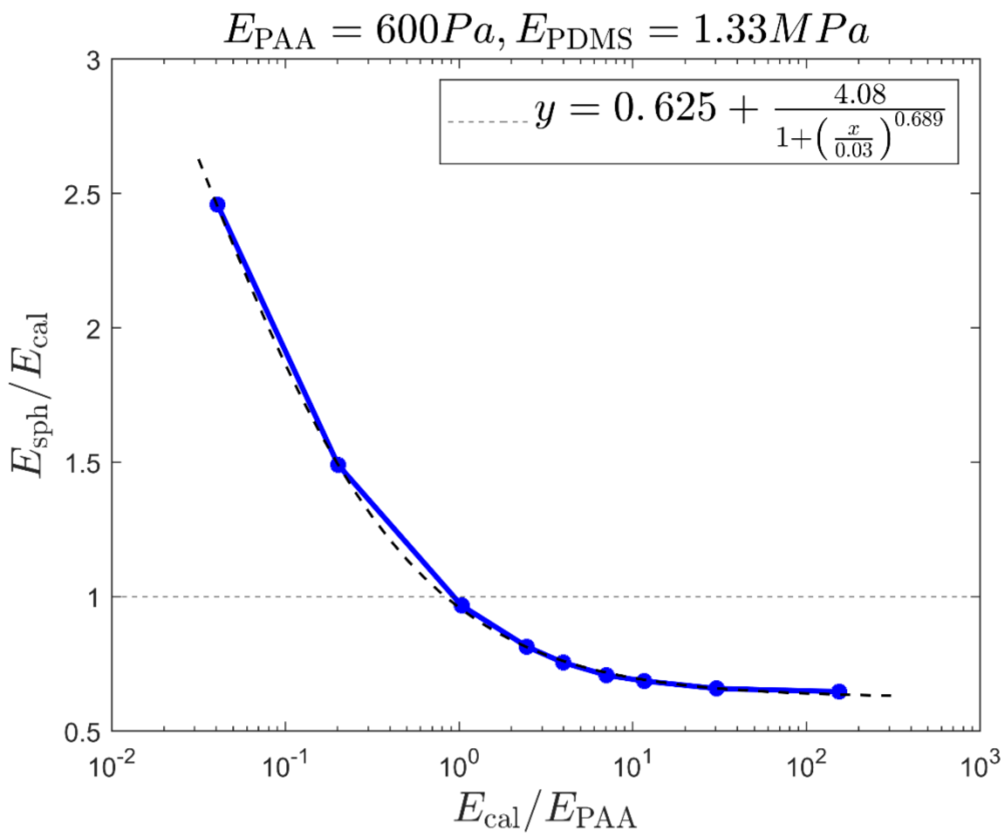


**Figure S15: Ratio of the calculated modulus $E_{cal}$ and the reference value (FEA input) of spheroid $E_{sph}$ as a function of the modulus ratio $E_{cal}/E_{PAA}$.** The relative difference is observed to be bounded as the spheroid becomes stiffer. A fitting function is proposed, and the calculated modulus can be corrected using it. Note that although the simulation is conducted with $E_{PAA} = 600Pa$, it is the modulus ratio $E_{sph}/E_{PAA}$ that matters.

Figure S15 shows the ratio of $E_{sph}$ and the input value $E_{cal}$ as a function of the modulus ratio $E_{cal}/E_{PAA}$. A fitting function is proposed here,

$$y = 0.625 + \frac{4.08}{1 + \left(\frac{x}{0.03}\right)^{0.689}} \qquad \text{(Eq. S3)}$$

We can apply this fitting function to correct the calculated modulus by,

$$E_{sph} = E_{cal}\left[0.625 + \frac{4.08}{1 + \left(\frac{E_{cal}}{0.03E_{PAA}}\right)^{0.689}}\right] \qquad \text{(Eq. S4)}$$

In our experiments, this ratio $E_{\mathrm{cal}}/E_{\mathrm{PAA}}$ lies within the range [1,10]. In this range, the calculated modulus overestimates the true modulus of the spheroid, with the overestimation bounded by 30%. Some correction factors $E_{sph}/E_{\mathrm{cal}}$ are provided in the table below.

| $E_{\mathrm{cal}}/E_{\mathrm{PAA}}$ | 1 | 2 | 3 | 5 | 8 | 10 |
|---|---|---|---|---|---|---|
| $E_{sph}/E_{\mathrm{cal}}$ | 0.9594 | 0.8391 | 0.7890 | 0.7417 | 0.7101 | 0.6982 |

Table S1: Correction factors $E_{sph}/E_{\mathrm{cal}}$ for given modulus ratios $E_{\mathrm{cal}}/E_{\mathrm{PAA}}$.

#### 3. Validation of PAA gel modulus measurements using both indentation method and parallel plate rheometer

To validate the accuracy of our indentation-based modulus measurements of the polyacrylamide (PAA) force-sensing layer used in the microrheometer, we performed independent mechanical testing using a commercial shear rheometer (TA Instruments DHR3). Shear stress was plotted as a function of shear strain for three PAA gel samples (Figure S16B), yielding linear fits with slopes corresponding to the shear modulus G:

Trial 1: G = 243.6 Pa

Trial 2: G = 248.8 Pa

Trial 3: G = 238.3 Pa

To convert shear modulus to Young's modulus, we assumed the PAA gel behaves as a nearly incompressible isotropic material with Poisson's ratio $\nu = 0.5$. Under this assumption, the relation $E = 2G(1+\nu) = 3G$ was used. This yielded a rheometer-based Young's modulus of 730.7 ± 15.9 Pa. This value falls within the range of our indentation-based measurements (638.68 ± 41.5 Pa (n = 18)); Fig. S16A), supporting the validity of our indentation method used in our microrheometer. These rheological measurements also confirmed that the PAA gel behaves as linear elastic material within the relevant strain range (up to 100%).

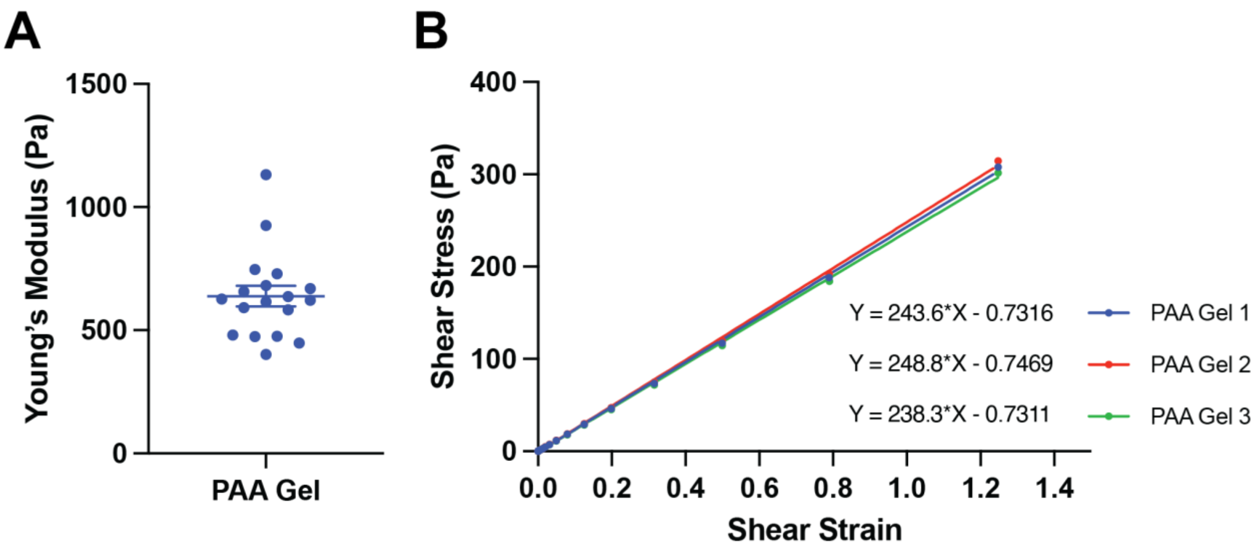

**Figure S16: Comparison of Young's modulus measurements of the Polyacrylamide gel using indentation method and a parallel plate rheometer.** A. Young's modulus of the PAA gel measured using the indentation method in the microrheometer across 18 independent measurements. The average modulus was 638.68 ± 41.5 Pa. B. Shear stress vs. shear strain curves measured using a TA Instruments DHR3 rheometer for three independently prepared gel samples. Linear regression fits are shown with corresponding slope values (shear modulus G). The measured average Young's modulus is 730.7 ± 15.9 Pa.

**References**[1] R. Long, M. S. Hall, M. Wu and C. Y. Hui, *Biophys* J, 2011, **101**, 643-650.

**Supplementary Movies**

**SMovie 1:** Piston movement in the Microrheometer under 10 kPa, T = 2 sec square pressure waves. This movie demonstrates the stability and responsiveness of the piston during dynamic loading.

**SMovie 2:** Piston movement in the Microrheometer under 10 kPa, T = 4 sec square pressure waves. This movie demonstrates the stability and responsiveness of the piston during dynamic loading.

**SMovie 3:** Piston movement in the Microrheometer under 10 kPa, T = 20 sec square pressure waves. This movie demonstrates the stability and responsiveness of the piston during dynamic loading.

**SMovie 4:** Compression of an MCF-10A spheroid under 15 kPa, T = 20 sec sinusoidal wave pressure input. This movie illustrates the spheroid's deformation dynamics during sinusoidal cyclic loading. Image dimensions: 433.44 x 330.24 µm

**SMovie 5:** Compression of an MCF-10A spheroid under 15 kPa, T = 20 sec triangular wave pressure input. This movie illustrates the spheroid's deformation dynamics during triangular cyclic loading.

**SMovie 6:** Compression of an MCF-10A spheroid under 15 kPa, T = 20 sec square wave pressure input. This movie illustrates the spheroid's deformation dynamics during square cyclic loading.

**SMovie 7:** Cellular dynamics of an MDA-MB-231 spheroids under 30% Δh/h compression over 24 hours. Images were taken every 5 minutes to visualize time-resolved cellular behavior during sustained compression. Image dimensions: 433.44 x 330.24 µm

**SMovie 8:** Cellular dynamics of an MDA-MB-231 spheroids under no compression over 24 hours. Images were taken every 5 minutes to visualize time-resolved cellular behavior in the absence of mechanical stress.

**SMovie 9:** Cellular dynamics of an MCF-10A spheroids under 30% Δh/h compression over 24 hours. Images were taken every 5 minutes to visualize time-resolved cellular behavior during sustained compression.

**SMovie 10:** Cellular dynamics of an MCF-10A spheroids under no compression over 24 hours. Images were taken every 5 minutes to visualize time-resolved cellular behavior in the absence of mechanical stress.

**SMovie 11:** Rotational behavior of compressed MCF-10A spheroids. PIV vector field is overlaid to visualize collective cell motion during compression. Image dimensions: 433.44 x 330.24 μm

**SMovie 12:** Rotational behavior of uncompressed MCF-10A spheroids. PIV vector field is overlaid to visualize collective cell motion in the absence of compression.
